# Prostate Cancer Reshapes the Secreted and Extracellular Vesicle Urinary Proteomes

**DOI:** 10.1101/2023.07.23.550214

**Authors:** Amanda Khoo, Meinusha Govindarajan, Zhuyu Qiu, Lydia Y. Liu, Vladimir Ignatchenko, Matthew Waas, Andrew Macklin, Alexander Keszei, Brian P. Main, Lifang Yang, Raymond S. Lance, Michelle R. Downes, O. John Semmes, Danny Vesprini, Stanley K. Liu, Julius O. Nyalwidhe, Paul C. Boutros, Thomas Kislinger

## Abstract

Urine is a complex biofluid that reflects both overall physiologic state and the state of the genitourinary tissues through which it passes. It contains both secreted proteins and proteins encapsulated in tissue-derived extracellular vesicles (EVs). To understand the population variability and clinical utility of urine, we quantified the secreted and EV proteomes from 190 men, including a subset with prostate cancer. We demonstrate that a simple protocol enriches prostatic proteins in urine. Secreted and EV proteins arise from different subcellular compartments. Urinary EVs are faithful surrogates of tissue proteomes, but secreted proteins in urine or cell line EVs are not. The urinary proteome is longitudinally stable over several years. It can accurately and non-invasively distinguish malignant from benign prostatic lesions, and can risk-stratify prostate tumors. This resource quantifies the complexity of the urinary proteome, and reveals the synergistic value of secreted and EV proteomes for translational and biomarker studies.

## INTRODUCTION

Human urine is produced when blood is filtered by the kidneys. This filtration retains proteins and nutrients, removes undesired metabolites and regulates pH and water levels. Excreted urine is less than 1% of pre-filtered volume^1^. After filtration, urine is stored in the bladder for minutes to hours before being voided through the urethra. In males, the urethra runs through the prostate, a gland that produces prostatic fluid^2^. Because urine can spend significant residence time within the genitourinary tract, it accumulates bioanalytes that reflect the current state of those tissues^3^. The urine is therefore a remarkably complex biofluid, and provides a non-invasive snapshot of both organismal state and genitourinary tissues. Its molecular composition can vary across individuals, and within a single individual over time^4^. Urine has been widely proposed as a non-invasive longitudinal biomarker matrix^5,6^, and specific DNA, RNA or protein species have been identified in urine that can serve as biomarkers^7–10^.

Proteins enter the urine in two ways: leakage at the glomeruli of the kidney and throughout the urogenital tract. It is believed that the vast majority of the excreted urinary proteome derives from tissues of the genitourinary tract, rather than leakage from the kidneys. Urogenitary proteins can enter the urine *via* passive release through cell death, through active translocation and as part of secreted extracellular vesicles (EVs)^11,12^. EVs are nanosized particles with a lipid bilayer released by cells into the extracellular milieu. They vary dramatically in size, ranging from 30 to 2,000 nm in diameter, and are heterogeneous in their mechanisms of biogenesis, molecular composition and function^13^. EVs play a crucial role in both physiology and in the pathogenesis of diseases, including cancer^14^. EV and secreted proteomes are hypothesized to be context-driven and tissue-specific^15^, but their presence, population variability and disease-relevance in urine remains poorly characterized.

To fill this gap, we generated comprehensive urinary proteomic profiles from 190 treatment-naïve men with a range of benign and malignant conditions. We demonstrate a simple protocol that uses urine to directly sample prostate proteins. This allows us to identify the tissue and subcellular origins of urinary proteins and EVs, and to quantify how the urine proteome changes over time in specific individuals. Urinary EVs, but not those released from prostate cancer cell lines nor secreted urinary proteins, accurately reflect prostatic tissue. Prostate tumor-specific urinary proteins accurately distinguish men with and without prostate cancer, and risk-stratify those already with the disease. Canonical EV markers are not effective in urine, but we identify context-dependent urine EV cargo that accurately marks specific urinary EV populations.

## RESULTS

### Digital rectal examination enriches urine for prostate proteins

The urine proteome is believed to derive almost exclusively from tissues of the genitourinary tract: the kidney, bladder and (in men) prostate^11^. Perturbation of the prostate gland using digital rectal examination (DRE) enriches for prostate-specific RNAs in urine^16^. While the mechanism is unknown, it is thought to occur by expelling prostatic fluid into the urethra, where it can be collected as part of first-catch urine. DREs are routine, minimally-invasive physical manipulations performed millions of times annually by oncologists and primary care physicians, and thus might provide a simple approach to enrich prostate-derived proteins in urine.

We therefore collected matched pre- and post-DRE urines from ten men (**Figure 1A**) and applied differential ultracentrifugation to separate urine soluble proteins (uSP) from urinary extracellular vesicles (uEVs). We further isolated two uEV populations based on size^17^: one at 20,000 x *g* (termed uEV-P20) and the other at 150,000 x *g* (uEV-P150; **Figure S1A**). To determine if a DRE influenced biophysical characteristics of EVs, we quantified EV diameter, number and morphology by nanoparticle tracking and by transmission electron microscopy (**Figure 1B** and **Figures S1B-D**). uEV biophysical characteristics were unchanged before or after a DRE (**Figures S1B-D**).

**Figure 1.**
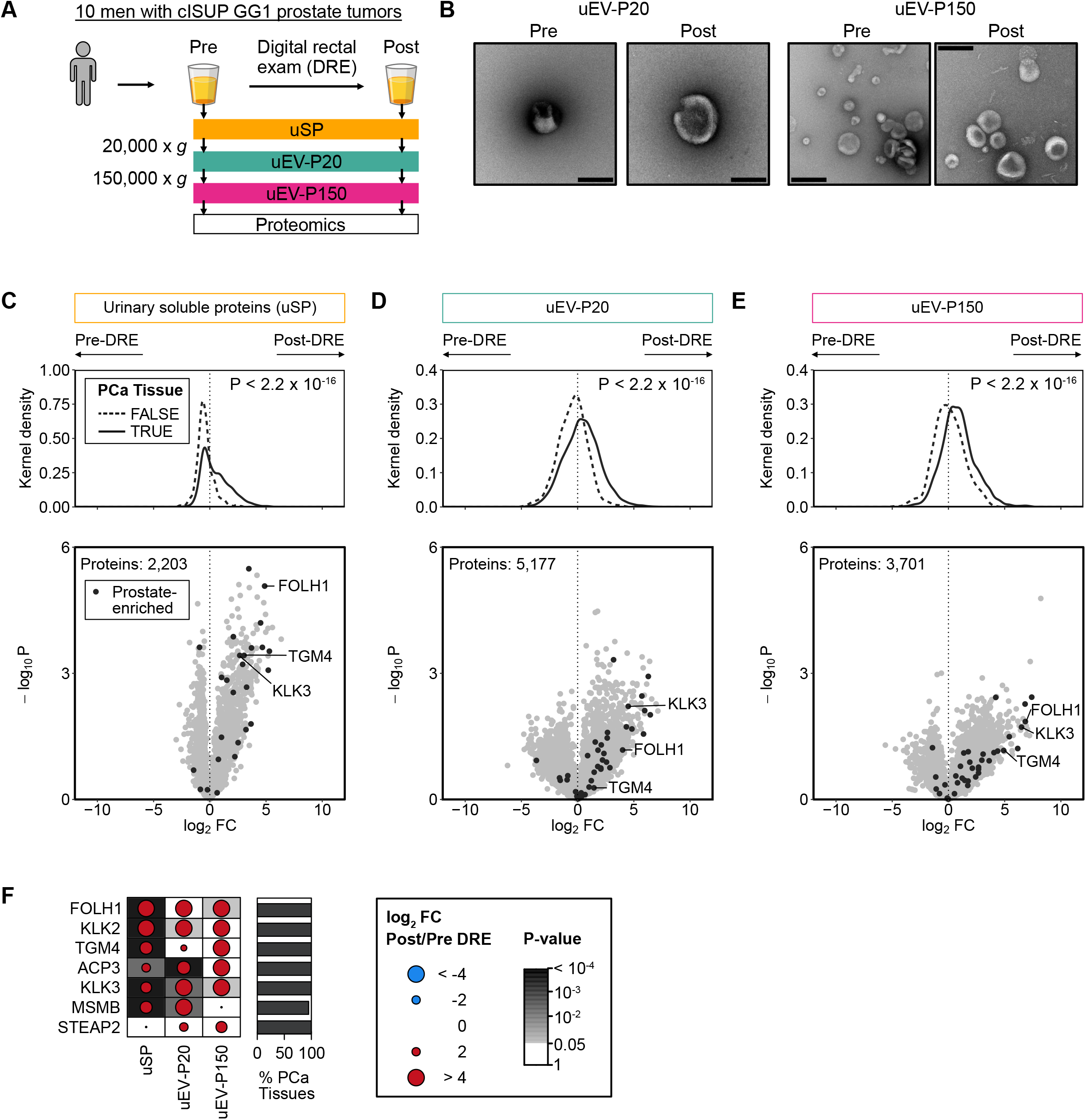
Digital rectal exam enriches for prostate tissue-derived proteins. **(A)** Matched pre- and post-DRE urine proteomes consisting of unfractionated urine (soluble proteins, uSP) and two subtypes of urinary extracellular vesicles (uEV) isolated by differential ultracentrifugation at 20,000 x *g* (uEV-P20) and 150,000 x *g* (uEV-P150). **(B)** Transmission electron microscopy images of uEVs isolated from pre- and post-DRE urine from a single individual. Scale bar: 200 nm. **(C-E)** Proteomic differences in pre- and post-DRE urines in **(C)** uSP, **(D)** uEV-P20 and **(E)** uEV-P150 fractions. Top panel: log_2_FC in protein abundance in urine, grouped by protein detection in prostate tissues^18,19^. Bonferroni-corrected P-values from Mann-Whitney U tests. Bottom panel: log_2_FC in protein abundances. Prostate-specific proteins as per the Human Protein Atlas are in black. **(F)** Differences in pre-*vs.* post-DRE urine (log_2_FC) for select prostate tissue-specific proteins^20,53,54^. Percentage of 157 prostate cancer tissues^18,19^ in which each protein was detected on the left. Background shading denotes P-value < 0.05 from a Wilcoxon signed-rank test. See also **Figure S1**.

To evaluate if a DRE increased the abundance of prostate-derived proteins, we measured the proteomes of each urine fraction using mass spectrometry. Prostate tissue-derived proteins were curated from three independent tissue proteomics datasets^18–20^, then annotated in urine. While the total number of urine proteins was unchanged by a DRE (4,064 ± 604 pre-DRE *vs.* 4,362 ± 511 post-DRE; P = 0.31; Wilcoxon signed-rank test), prostate tissue-derived proteins were significantly more abundant in post-DRE urine (P < 2.2 x 10^−16^; **Figures 1C-E**). These included classic prostate marker proteins like PSMA (*FOLH1*) and PSA (*KLK3*)^20^ (**Figures 1F and Figure S1E**). Thus, a DRE significantly enriches the urine for proteins of prostate origin but does not influence EV biophysical characteristics, suggesting the latter may be relatively tissue-independent.

### The subcellular origins of urinary proteins

To investigate urinary protein heterogeneity, we next collected post-DRE urines from 190 men: 64 men with no cancer diagnosis and 126 with untreated prostate cancers. The prostate cancer patients reflected the full risk spectrum of primary disease, with biopsy ISUP Grade Groups (GG) ranging from low (GG1) to high (GG5; **Figure 2A**). We isolated uSP, uEV-P20 and uEV-P150 from post-DRE urine and quantified their proteomes (**Figure 2B**; **Table S1**). uEV biophysics and urine protein counts were largely independent of disease status, age or serum PSA levels (**Figures S2A-G**). The uEV-P20 and uEV-P150 fractions were biophysically similar in size, morphology and particle count (**Figures 2C-E**).

**Figure 2.**
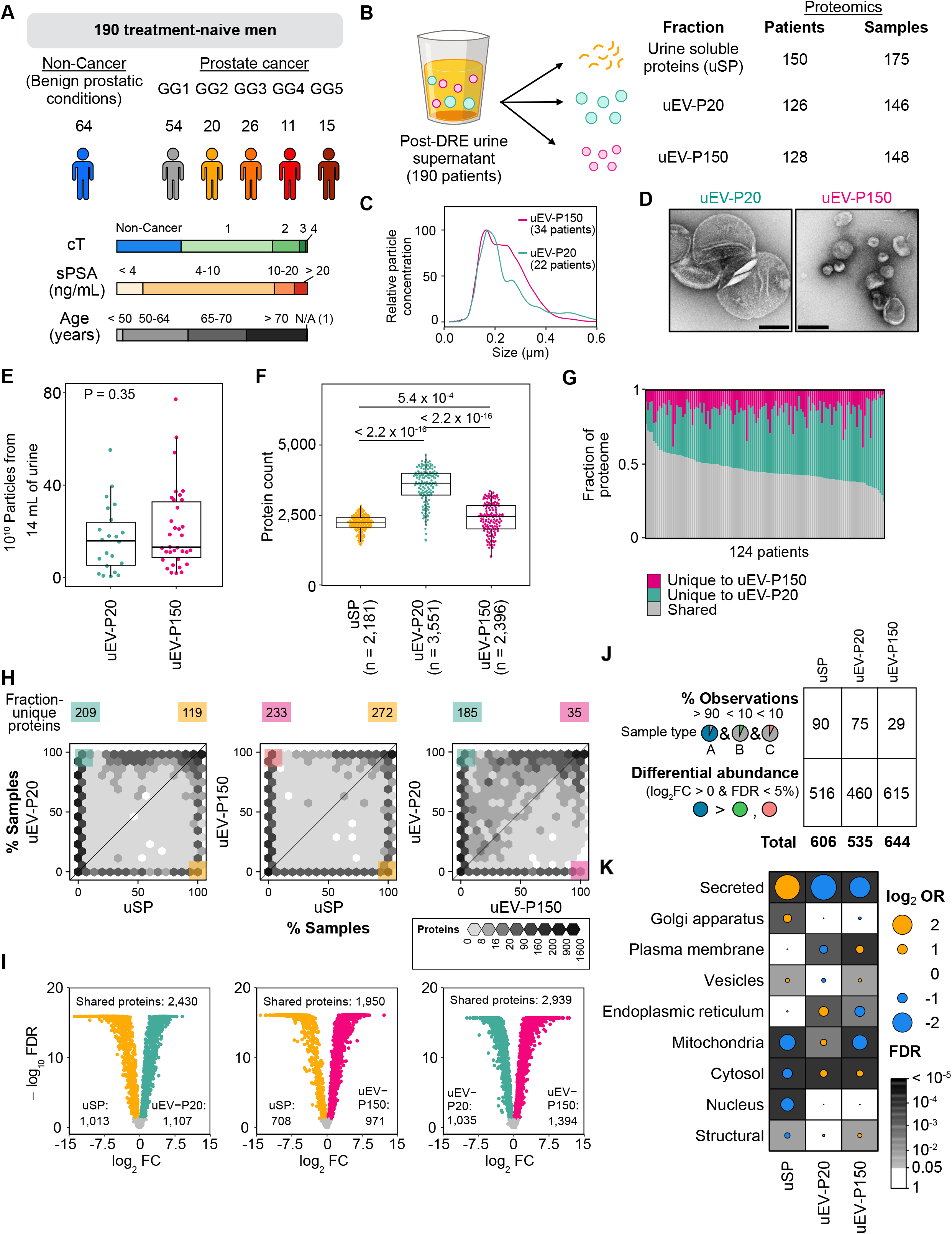
Post-DRE uEV fractions harbor distinct protein cargo. **(A)** Cohort overview. Clinical (biopsy-based) International Society of Urological Pathology Grade Group (GG); cT: clinical T category; sPSA: Serum Prostate Specific Antigen (ng/mL); Age: Age at diagnosis (years). **(B)** Post-DRE urine fractions analyzed by proteomics, total patient number and total sample number per fraction. **(C)** Distribution of particle sizes determined by nanoparticle tracking analysis as mean of n = 22 of uEV-P20 and n = 34 of uEV-P150. **(D)** Representative transmission electron microscopy images of uEV fractions from one cISUP Grade Group 1 patient. Scale bar: 200 nm. **(E)** Particle concentration for 22 uEV-P20 and 34 uEV-P150. P-value from Mann-Whitney U test. **(F)** Number of proteins quantified by mass spectrometry. Dots represents samples (Samples: 175 uSP, 146 uEV-P20, 148 uEV-P150). P-values from Mann-Whitney U tests. **(G)** Fraction of uEV-P20 and uEV-P150-unique proteins from 124 patients with matched uEV fractions. **(H)** Number of samples each protein was detected in (n = 86 patients with all three fractions). For each pairwise-comparison, the numbers of proteins present in > 90% of samples in one sample type and < 10% of the other are labeled on top of each panel. **(I)** Differences in shared protein abundance between fractions. Significant differences (FDR < 0.05, Wilcoxon signed-rank test) are in green, pink or yellow. Total differentially abundant proteins are in bottom corners. **(J)** Fraction-enriched proteins either unique to one fraction or differentially abundant in one fraction relative to the other two. **(K)** Odds ratio of gene set enrichment for each subcellular localization^55^ for proteins from Figure 2J. Grey background shading indicates FDR < 0.05 (Fisher’s Exact Test). See also Figures S2 and S3.

Despite these biophysical similarities, the proteomes of different urine fractions were significantly different. Many urine EV proteins were identified as EV cargo in previous studies (**Figures S3A-B**)^17,21–24^. The uEV-P20 fraction had the most detectable proteins (**Figure 2F**) and was the most biophysically and proteomically diverse, suggesting heterogeneity in vesicular type or origin (**Figure 2G**). uEV fractions were more similar to one another than to the soluble protein fraction, consistent with the detected EV proteins being true cargo rather than co-isolated urinary proteins (**Figures S3C-E**).

To determine the differential origins and biology represented by each fraction, we performed differential proteome analysis. We identified proteins detected in one fraction but not the others (**Figure 2H**) and proteins detected at different abundances across fractions (**Figure 2I**). Each fraction was defined by presence of ∼50 proteins and differential abundance of ∼500 others (**Figure 2J**). Fraction-specific proteins tended to arise from specific subcellular compartments. Urine soluble proteins were typically secreted or derived from the Golgi apparatus, while uEV-P20 proteins derived from mitochondria or endoplasmic reticulum and uEV-P150 proteins from the plasma membrane (**Figure 2K**).

### uEVs but not cEVs are reflective of the prostate tissue proteome

We next quantified how well each of the three urine fraction proteomes reflects the proteome of prostate tissue^18,19^. A majority (67%) of all proteins detected in prostate tissue were detected in one or more of the three urine fractions (**Figure 3A**). EV fractions were a much richer source of prostate-derived proteins than urine soluble proteins. Only 116 prostate-derived proteins were identified in unfractionated urine, whereas 2,439 were identified only in one or both of EV fractions and 4,968 in both EV and non-EV urine. Protein abundances were well-correlated between urinary and tissue proteomes (**Figure 3B**), with uEV-P20 being the best surrogate for prostate tissue.

**Figure 3.**
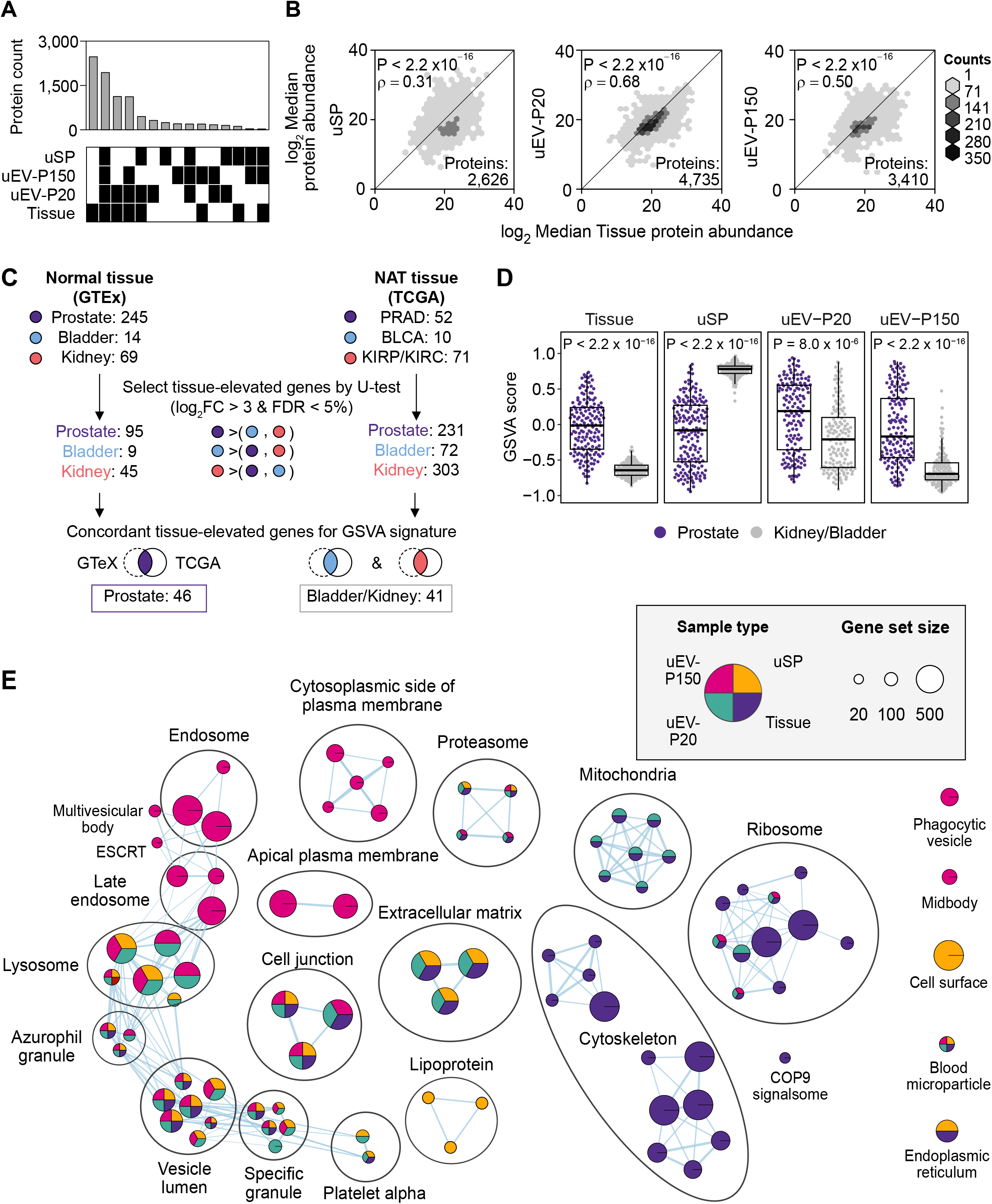
uEV proteome closely reflects the prostate tissue proteome. **(A)** Overlap in proteins quantified in each sample type. Samples: Soluble proteins (uSP) = 175, uEV-P150 = 148, uEV-P20 = 146, Tissue = 157; Proteins: uSP = 3,150, uEV-P150 = 3,878, uEV-P20 = 5,462, Tissue = 7,438. **(B)** log_2_ median protein abundance between prostate tissue and uSP (left), uEV-P20 (middle) and uEV-P150 (right). Spearman’s rank correlation and its p-value are shown. **(C)** Analysis strategy to identify tissue-associated genes in RNA-seq of normal or NAT prostate, bladder and kidney from GTEx^25^ or TCGA^26–29^ (see **Table S4**). **(D)** Gene set variation analysis scores of sample types based on tissue-specific signatures. P-values from Mann-Whitney U test. **(E)** GO:CC gene sets over-represented in each sample type (see also **Table S3**). Only significant gene sets (FDR-adjusted hypergeometric p-value < 0.05) are shown.

To further quantify the tissue provenance of urinary proteins, we used RNA-seq data from normal tissues (GTEx^25^) and from normal tissue adjacent to tumors (NAT, TCGA^26–29^) to identify transcripts enriched in prostate, kidney or bladder (**Figure 3C**). Differentially abundant transcripts were then used as signatures of tissue origin to quantify the contribution of each tissue to each sample. In uEVs, prostate proteins were very significantly more abundant than non-prostate proteins (P < 1 x 10^-5^). In contrast, the soluble urine fraction (uSP) showed the inverse trend: it was significantly depleted in prostate-derived proteins (P < 2.2 x 10^-16^; **Figure 3D**). The soluble protein fraction (uSP) was enriched in functions classically associated with blood (lipoprotein, blood microparticle; **Figure 3E**). The soluble proteome was also highly enriched in cell surface proteins, likely from the shedding of extracellular domains^30,31^. Proteins involved in multivesicular body biogenesis were over-represented in uEV-P150, suggesting an enrichment of exosomes^32^. Consistent with the univariate protein analysis, uEV-P20 fraction more closely reflected prostate tissue, harboring proteins originating from mitochondria, ribosome and extracellular matrix^33^. These data suggest that the prostate primarily sheds EVs into urine, and that these can provide a robust non-invasive proxy of its proteome.

Cell line conditioned media has been widely used to study secreted cellular components^34,35^. To ascertain if cell line-derived EVs (cEVs) accurately reflect tumor tissue, we isolated and proteomically characterized cEVs from five prostate cell lines (**Figure 4A**). EVs from urine and from cell line conditioned media displayed similar biophysical characteristics (**Figures S4A-D**). More proteins were detected in cEV-P20 than in cEV-P150, but fewer than in whole-cell lysates (**Figures S4E-F**). Cell line whole-cell lysates closely resembled tumor tissue in both protein composition (**Figure 4B**) and abundance (**Figure 4C**). By contrast uEVs and cEVs differed, particularly in abundance (**Figures 4D-E**). Urinary EVs were better surrogates for prostate tissue than were EVs derived from cell lines (**Figure S4G**). Commonly used EV markers like CD9, CD81 and CD63^36,37^ were amongst the proteins more abundant in EV-P150 than EV-P20 in both cell lines and urines (**Figure 4F** and **Figure S4H**), while other EV markers like FLOT1 were discordant between cell lines and urine (**Figure 4F**). Mitochondrial proteins were over-represented in cEV-P20 and plasma membrane proteins in cEV-P150 (**Figure 4G**). Thus, post-DRE EV-associated proteins more accurately reflect prostate tissue than prostate cancer cell line EVs or, particularly, post-DRE urine soluble proteins.

**Figure 4.**
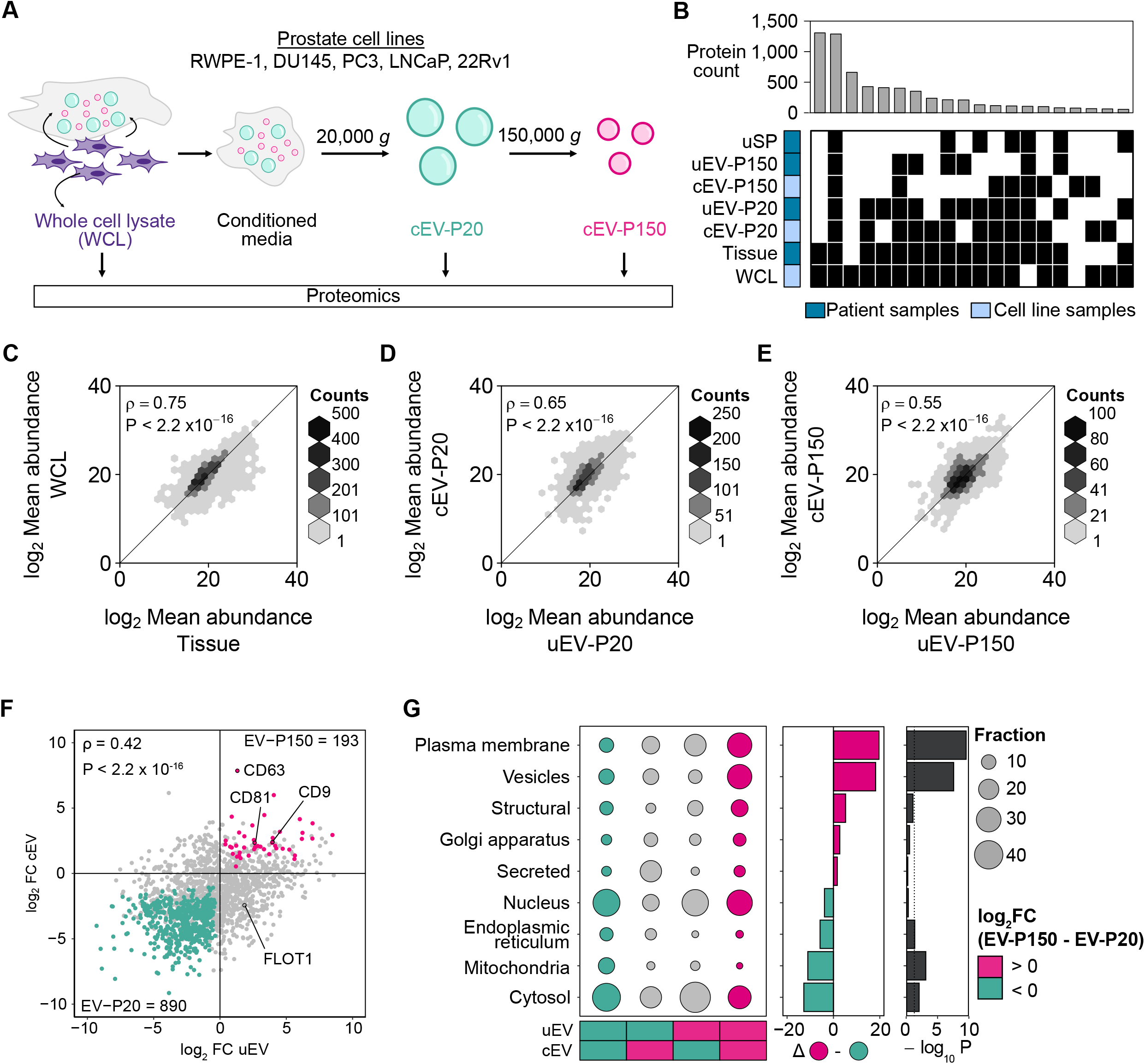
Prostate cancer cell line EVs do not fully reflect prostate fluid EVs. **(A)** Overview of cell line EV (cEV) isolation from conditioned media. **(B)** Overlap in proteins quantified between sample types. **(C-E)** Correlation between log_2_ mean protein abundance in patient and cell line fractions. Cell line protein abundances are mean of three processing replicates from five cell lines. (F) log_2_FC between EV-P150 and EV-P20 in uEV (x-axis) *vs.* cEV (y-axis). Significantly differentially abundant proteins in both uEV and cEV (Mann-Whitney U test < 0.05) are pink (higher in EV-P150) or green (higher in EV-P20). (G) Organellar enrichment for proteins (two-proportion z-test) in each quadrant from Figure 2D (left panel), annotated with differences in fraction (middle panel) between EV-P150 (top right quadrant) and EV-P20 (bottom left quadrant) and P-value (right panel). See also **Figure S4**.

### Biomarker potential of urinary proteomes

For a biomarker to be useful, it needs to be robust to a variety of types of errors. While urine is not prone to spatial biases, it is unclear to what extent an individual’s urine proteome is stable over time. To quantify the temporal stability of post-DRE urine, we evaluated longitudinal samples in five prostate cancer patients over several years. From each patient, we collected post-DRE urine at multiple timepoints; all patients were managed by active surveillance without any indication of clinical progression (**Figure 5A**; **Table S1**). We used variance analysis to quantify which proteins were more variable between samples of a single patient (intra) or across individuals (inter). On average, proteomes were more similar within patients than between them, for both EV cargo and urine-soluble proteins (**Figure 5B**). To identify proteins that might be particularly useful as biomarkers, we identified those that were longitudinally stable using the intraclass correlation coefficient (ICC)^38^. The higher a protein’s ICC, the less its variance in protein abundance is caused by random fluctuations over time. A subset of proteins was highly longitudinally stable in each fraction (**Figure S5A**), and these comprise excellent candidate biomarkers because they are robust to physiological variability over several years.

**Figure 5.**
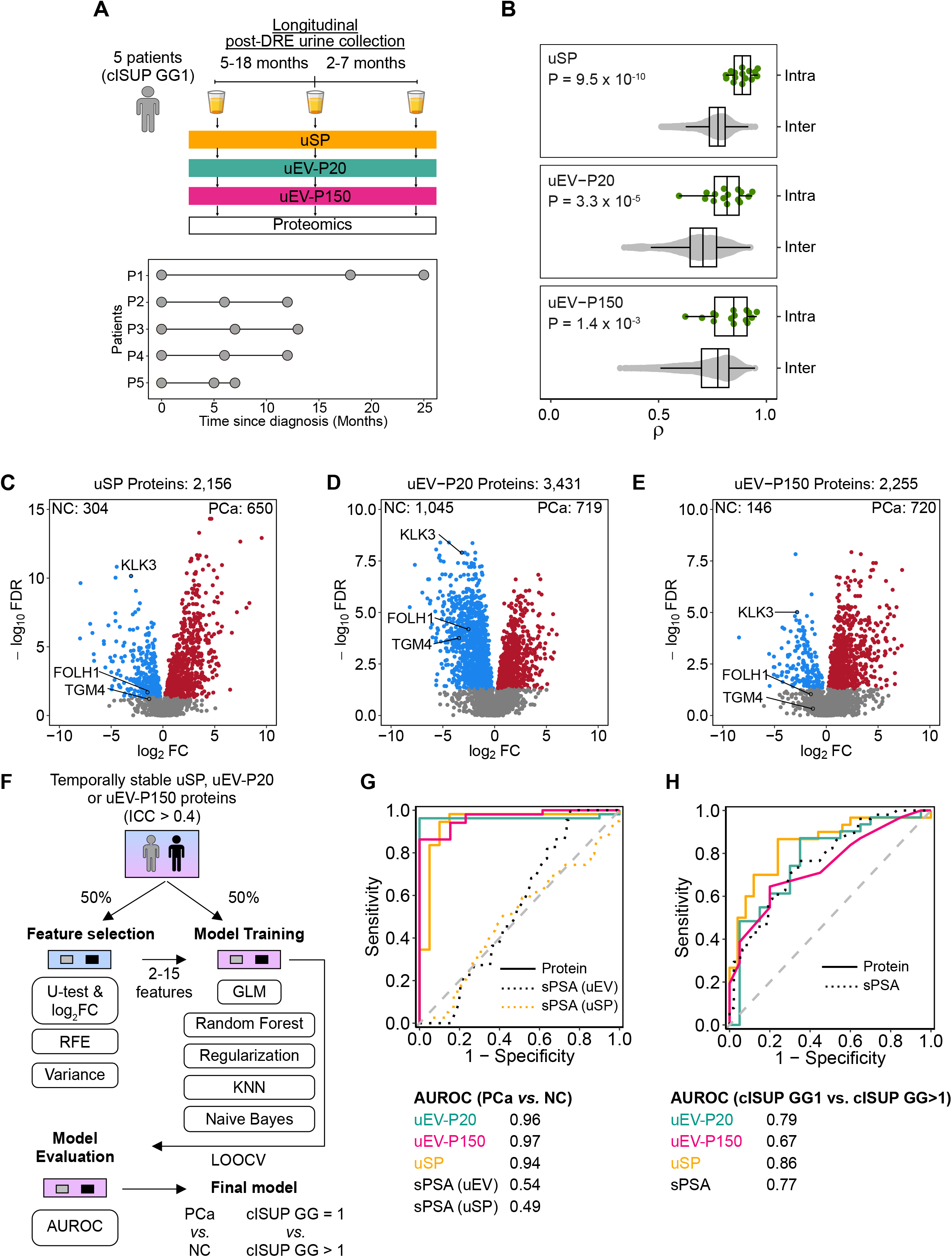
uEV proteome is temporally stable and reflects clinical behavior. **(A)** Timepoints for each post-DRE urine collection in a cohort of five patients with cISUP GG1 tumors on active surveillance that did not upgrade over the course of urine collection. **(B)** Protein abundance (Spearman’s ρ) within individuals (Intra, green, 5 patients with cISUP GG1 tumors), and between individuals (Inter, grey; patients: uSP = 150, uEV-P20 = 126, uEV-P150 = 128). P-values from Mann-Whitney U tests. **(C-E)** Significantly differentially abundant proteins between prostate cancers (PCa, red) *vs.* non-cancers (NC, blue) using a Mann-Whitney U test FDR < 0.05. NC patients for uSP and uEV are not matched. Patients: uSP_NC_ = 39, uSP_PCa_= 136, uEV-P20_NC_ = 22, uEV-P20_PCa_ = 132, uEV-P150_NC_ = 25, uEV-P150_PCa_= 131. **(F)** Strategy for building models to classifying PCa *vs.* NC, and cISUP GG1 *vs.* GG>1. **(G-H)** ROC curves for multi-protein models (solid line) and serum PSA (dotted line) in classifying **(G)** prostate cancer *vs.* non-cancer, and **(H)** cISUP GG1 *vs.* cISUP GG>1. See also **Figure S5**.

Next, we sought to evaluate the biomarker potential of urine-soluble and uEV proteins for predicting prostatic disease. In our cohort, patients with and without prostate cancer had similar serum PSA abundances (P = 0.34; **Table S1**). Despite this similarity, thousands of soluble (**Figure 5C**) and EV proteins (**Figures 5D-E**) differed between patients with and without cancer. Intriguingly, PSA (*KLK3*) protein abundance was consistently significantly reduced in all fractions of urine from prostate cancer patients relative to men without a cancer diagnosis^2^, despite being increased in serum and in tumor regions^18^. The specific differentially abundant proteins varied between fractions (**Figures S5B-C**).

To create biomarkers of prostatic disease, we focused on proteins that were longitudinally stable (ICC > 0.4), which retained 806 uSP proteins, 1,322 uEV-P20 proteins and 674 uEV-P150 proteins. We used statistical machine learning to create and validate classifiers independently for each urine fraction (**Figure 5F**). First, we created classifiers that distinguish cancer from non-cancer based solely on urine proteins; these had AUCs ranging from 0.94-0.97, significantly outperforming serum PSA (**Figure 5G**). Next, we applied the same methodology to the more challenging task of distinguishing low-grade from high-grade cancer. This is an important clinical question, as low-grade disease is typically managed by active surveillance and higher-grade disease by definitive local therapy. Urine distinguished low-from high-grade prostate cancer with AUCs ranging from 0.67 to 0.86 (**Figure 5H**), again matching or exceeding serum PSA. Intriguingly, signatures of grade also informed on disease status, reflecting an overlap in determinants of initiation and progression, as seen in studies of prostate cancer genetic drivers^39^ (**Figure S5D**). Proteins within these signatures exhibit longitudinal stability and were univariately associated with disease status (**Figure S5E**). These data suggest that the urine proteome is an untapped source of biomarkers for genitourinary disease.

### Markers of prostate tumor uEVs

Post-DRE urine can non-invasively sample the prostate tissue proteome and has significant biomarker potential. However, the soluble fraction and the protein cargo of different size EVs differ in their origin and association with disease phenotypes. As examples, SPOCK1 protein was significantly lower in the soluble fraction of urine from cancer patients but not in uEVs, while HLA-DRB5 shows the inverse (**Figure S5E**). This interplay between subcellular origin, clinical phenotype and urine fraction is summarized in **Figure 6A**, and suggests that protein cargo is selectively packaged into EVs in processes that are dysregulated during malignant transformation (**Figure S6A**).

**Figure 6.**
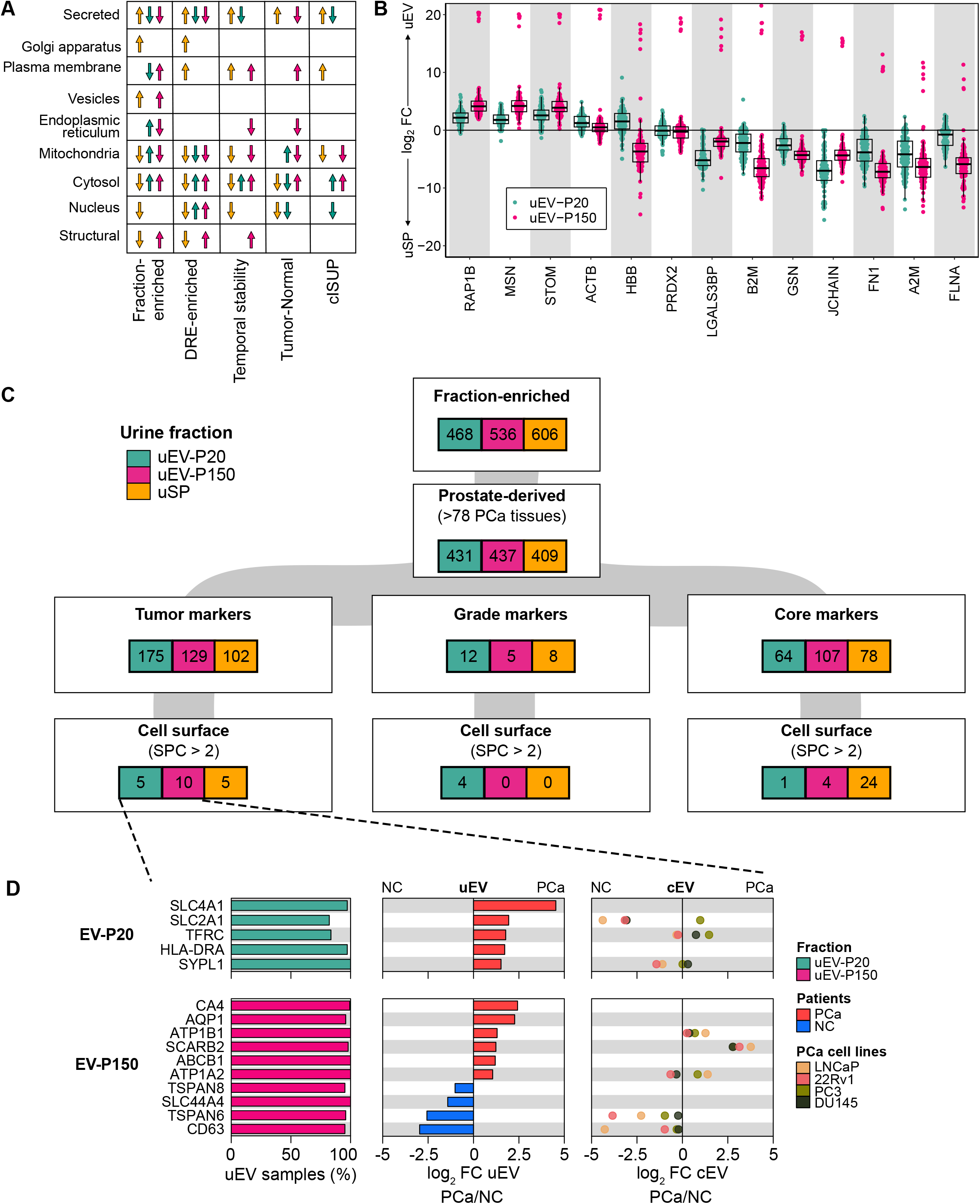
EV cargo is context-dependent. **(A)** Proteins differentially abundant in each condition and urine fraction by compartment. **(B)** Differences in protein abundance of extracellular vesicle markers from^40^ in matched uEV-P20 (green) and uEV-P150 (pink) relative to uSP. **(C)** Fraction-specific proteins were determined by differential protein abundance and frequency of detection (Figure 2J). Proteins in each subset were filtered based on the following criteria. Tumor markers: |log_2_FC|_PCa/NC_ > 1 & FDR _PCa/NC_ < 0.05; Grade markers: |log_2_FC|_cISUP>1/cISUP=1_ > 1 & unadjusted P _cISUP>1/cISUP=1_ < 0.05; Core markers: |log_2_FC|_PCa/NC_ < 1 & FDR _PCa/NC_ > 0.05 & |log_2_FC|_cISUP>1/cISUP=1_ > 1 & unadjusted P_cISUP>1/cISUP=1_ > 0.05 & > 90% of samples & < 25% least variant proteins. Tumor, grade and core markers were further filtered to select cell surface markers (SPC^41^ > 2). PCa: Prostate cancer; NC: Non-cancer; SPC: Surface prediction consensus^41^. P-values from Mann-Whitney U test. **(D)** Tumor markers with predicted cell surface localization from Figure 6C. Proteins are annotated with the frequency of detection in each fraction (left panel), and differential abundance in PCa *vs.* NC in uEV (middle panel) and cEV (right panel). See also **Figure S6**.

This heterogeneity in origins and association with clinico-epidemiologic characteristics highlights the importance of accurately isolating specific EV populations. The gold-standard intensive centrifugation used here is expensive and time-consuming, so affinity-based isolation methods are strongly preferred for rapid translational and clinical studies. Affinity methods require identification of proteins characteristic of specific EV subpopulations^40^. We evaluated the performance of the 13 best available EV markers^40^. Four of these were elevated in uEV-P150 relative to soluble urine proteins and five were elevated in uEV-P20. Others were depleted in uEVs, and none showed the large effect sizes or small inter-sample variability needed for ideal affinity-based markers (**Figure 6B**). Thus canonical EV protein markers do not appear to be optimal for uEV identification and isolation.

We therefore sought to identify protein markers to distinguish prostate-derived uEVs. We selected proteins that were both distinct to one urine fraction (**Figure 2J**) and are known to be present in prostate tissue. We separated these into three protein subsets: those associated with presence of a tumor, those associated with tumor grade and disease-invariant “core” proteins (**Figure 6C**). These subsets were functionally distinct, with prostate-specific secreted proteins such as KLK3 and ACPP (Cluster C_uSP_8) being specific to the urine soluble protein fraction but not uEVs, and GTP binding proteins specific to the uEV-P150 fraction (Cluster C_P150_8)^15^ (**Figures S6B-D**).

Finally, to identify specific actionable markers for uEV affinity studies, we selected predicted cell surface proteins from each subset^41^ (**Figure 6C** and **Figures S6E-F**). The resulting five uEV-P20 and ten uEV-P150 tumor markers that differed in patients with cancer and benign disease include the classical EV marker CD63^36,37^ (**Figure 6D**). Grade markers (**Figure S6E**) contain markers such as ITGB2 and SLC4A1 in the uEV-P20 fraction. Disease-invariant core markers of uEV-P150 include classical EV markers CD9 and CD81, which were frequently detected in uEV-P150 (**Figure S6F**). These data indicate that the EV proteome is context-specific (**Figure S6G**), and that protocols for rapid, specific isolation of EV subpopulations may differ from those useful in plasma or some other tissues.

## DISCUSSION

Urine contains a complex mixture of proteins that differ in their form of release and tissue of origin, resulting in a dynamic range of concentrations spanning ten orders of magnitude^4^. To better define the urinary proteome, we used fractionation to distinguish urine-soluble proteins from proteins carried in urinary extracellular vesicles (uEVs). uEVs of different sizes and densities contained proteins from different subcellular origins, suggesting distinct biogenesis^33,42^. uEVs, particularly the uEV-P20 population, appear to derive heavily from prostate tissue, and to accurately reflect its proteome. By contrast, neither the soluble urine proteome nor cell line-derived EVs did so. This highlights the need to prioritize patient-derived EVs in translational studies. This is particularly key for diseases where models that faithfully recapitulate aspects of the natural history like are lacking, such as hormone sensitivity and hypoxia in prostate cancer.

Prior work has rigorously quantified the role of factors such as sample collection, processing^2,43,44^ and storage^45^ on urine proteomes. We show that digital rectal examinations are a simple way to enrich for prostate proteins and EVs in first-catch urine. While post-DRE urine proteomes can vary over time^4,46^, a subset of specific proteins are temporally stable over many months and are well suited for non-invasive sampling. We found that the protein composition of urinary proteomes uniquely differs as a function of tumor grade, and when comparing prostate cancers to non-cancers. These differences in the secreted proteome between groups likely reflect altered signaling and intercellular communication by cancer cells, or as a compensatory mechanism for altered tumor metabolism or cellular stress^47,48^. Therefore, we show that EV cargo is context dependent and is associated with different cellular phenotypes, such as disease.

Further proteomic interrogation of urinary EVs in individuals of different ancestries, ages and in different clinical scenarios^49^ could offer new insights into genitourinary biology. However, the complexity of EV isolation by differential ultracentrifugation and requirement for high starting material can be limiting^17^. Affinity-based capture of specific EV subpopulations using cell surface markers is appealing^50^. Intriguingly, uEVs appears to be characterized by cell surface markers different from other EV subpopulations^40^. Thus new protocols to rapidly isolate EVs from urine, are urgently needed to maximize its utility for biomarker and other translational studies.

Since our unique dataset represents one of the largest urine proteomics studies, we were able to leverage longitudinally stable proteins to derive new biomarkers for prostatic diseases. Future validation using robust quantitation by targeted proteomics assays with stable isotope labeled standards^49,51^ in large, racially diverse cohorts will be required. The full data resource presented here is accessible *via* an interactive portal (http://kislingerlab.uhnres.utoronto.ca/ev/home/) to facilitate investigation into the urinary secreted and EV proteomes.

## METHOD DETAILS

### EV isolation from urine

Urinary extracellular vesicles (uEV) were isolated by differential ultracentrifugation as previously described ^17^. Briefly, 14 mL of frozen urine supernatant was thawed at 4°C, then diluted to a volume of 35 mL with isotonic buffer (250 mM sucrose, 10 mM HEPES, 1 mM EDTA, pH 7.4). The urine was centrifuged at 20,000 x *g* for 30 minutes at 4°C (k-factor 1,790) in an Optima XPN-80 ultracentrifuge (Beckman Coulter) equipped with a SW32 Ti swinging bucket rotor (R_min_ 67, R_max_ 153, Beckman Coulter) to pellet EVs. The 20,000 g pellet (P20) was treated with 500 mM of dithiothreitol (DTT) at 37°C for 30 min to reduce the uromodulin network, and centrifuged a second time at 20,000 x *g* for 30 min at room temperature. The P20 pellet was resuspended in 1mL of cold PBS, and centrifuged at 18,210 x *g* for 30 mins (Eppendorf Centrifuge 5430 R, FA-45-48-11 rotor, k-factor 198). The supernatant from the first and second centrifugation steps were combined and centrifuged at 150,000 x *g* for 2 h at 4°C (SW32TI swinging bucket rotor, k-factor 239) in an ultracentrifuge to pellet EVs. The 150,000 x *g* pellet (P150) was resuspended in high pH buffer then passed twice through a 0.22 µm filter. Samples were centrifuged again at 150,000 x *g* for 2 hours at 4°C to pellet uEV-P150. The P20 and P150 pellets containing uEV-P20 and uEV-P150, respectively, were resuspended in 100 µL of 50% *2,2,2*-trifluoroethanol in PBS, flash-frozen in liquid nitrogen, and stored at − 80°C until proteomics analysis.

### EV isolation from cell culture conditioned media for proteomics

All cell lines were grown to 70-80% confluency then washed three times with phosphate-buffered saline (PBS) and serum-starved for 48 hours prior collection of conditioned media. The cell line conditioned media containing EVs (cEV) was collected and centrifuged at 500 x *g* for 10 minutes then 2,000 x *g* at 4°C for 30 minutes to clear cell debris. The supernatant was concentrated to a volume of 4-5 mL (if using EVs for biophysical studies) or to a volume of 20-30 mL (if using EVs for proteomics) in a 100 kDa MWCO ultrafiltration concentrator (Millipore). EVs were isolated from conditioned media by differential ultracentrifugation in a SW32Ti swinging bucket rotor as described above. Conditioned media was topped off with PBS as required. Unlike the EV isolation protocol for urine described above, the first P20 pellet was not treated with DTT as cell line conditioned media is not expected to contain uromodulin. cEVs were collected from the 20,000 x *g* (cEV-P20) and 150,000 x *g* (cEV-P150) pellets.

### Urine proteomics

Proteomic profiles of the soluble protein fraction were generated from 250 µL of urine supernatant (following 2,000 x *g* centrifugation). Urine was prepared for proteomics using the MStern protocol^56^. For each sample, 2 pmol of *Saccharomyces cerevisiae invertase* was added as a sample preparation control. Proteins in each sample were reduced with 5 mM DTT and incubated for 30 min at 60°C. To prevent re-formation of disulfide bonds, 25 mM iodoacetamide was added and samples were incubated at room temperature for 30 min in the dark. Liquid in the following steps were passed through the MStern wells using vacuum suction unless otherwise stated. The membrane was equilibrated with 50 μL of 70% ethanol, then washed twice with 100 mM ammonium bicarbonate (ABC). Samples were added to the wells and passed through the membrane. Each well was washed twice with 100 μL of ABC to remove salts, then proteins were digested with 1 μg of mass spectrometry grade Trypsin/Lys-C enzyme mix (Promega) in 50 μL of digestion buffer (100 mM ABC, pH 8.0, 1 mM CaCl_2_, 5% acetonitrile). To ensure that the proteins are in contact with the digestion buffer, the digestion buffer was passed through the membrane by centrifugation, and the flow-through was reapplied on top of the membrane. Protein digestion was performed at 37°C for four hours. Samples were resuspended in the well by gentle pipetting every two hours. Peptides were collected by centrifugation, and remnant membrane-bound peptides were eluted with 50 μL of 50% acetonitrile and combined with the previous flow through. Samples were dried in a SpeedVac vacuum concentrator (Thermo). Dried peptides were resuspended in 0.1% trifluoroethanol in water and desalted using homemade solid phase extraction stage tips containing 3 plugs of 3M^TM^ Empore ^TM^ C18 membrane^57^. Peptides were quantified by NanoDrop (Thermo Scientific). 2 μg of peptides were loaded on column.

### EV proteomics

Cell line or urine EVs and cell pellets from whole cell lysates (WCL) were prepared for mass spectrometry as previously described^19^. Briefly, EVs or cells in 50% *2,2,2*-trifluoroethanol were lysed by freeze-thaw, then incubated at 60°C for 1 h to extract proteins. Then, proteins were reduced with 5 mM of DTT, alkylated with 25 mM of iodoacetamide, and digested overnight at 37°C with a 2 μg Trypsin/Lys-C enzyme mix (Promega). The next day, the enzymatic digest was quenched with 1% formic acid, and samples were desalted with homemade C18 StageTips (see above) prior to LC-MS analysis. iRT peptide standards (Biognosys) were spiked into reconstituted peptides at a 1:100 dilution according to manufacturer’s instructions.

### Mass spectrometry and data processing

Peptides were separated on a 50 cm C18 reverse phase EASY-Spray LC column interfaced with an EASY-nanoLC 1000 system over a 2-hour gradient (EVs and urine) or 4-hour gradient (WCL). Mass spectrometry for uEVs, cEVs, and WCL was performed on a Q-Exactive HF mass spectrometer coupled to an EASY-Spray ESI source (Thermo Scientific). Mass spectrometry for urines was performed on a Fusion Tribrid mass spectrometer coupled to an EASY-Spray ESI source (Thermo Scientific). All datasets were acquired in data-dependent acquisition mode. Raw files for each urine fraction were searched separately in MaxQuant^58^ (v.1.5.8.3 for uSP samples and v.1.6.2.3 for uEV samples) at a single site using a UniProt human protein sequence database (complete human proteome with isoforms). Searches were performed with a maximum of two missed cleavages, and carbamidomethylation of cysteine as a fixed modification. Variable modifications were set as oxidation of methionine. The false discovery rate for the target-decoy search was set to 1% for protein and peptide. Peptide detection was performed with an initial precursor and fragment mass deviation threshold of 10 and 20 parts per million respectively. Intensity-based absolute quantification (iBAQ), label-free quantitation, and match between runs (matching and alignment time windows set as 0.7 and 20 min respectively) were enabled. The *peptides.txt* output files from each MaxQuant search were parsed into an in-house database for protein grouping^59^. Protein abundances (gene-centric) were determined from peptide abundances using the iBAQ algorithm^60^ (**Table S2**). Reverse hits (false positives from target-decoy search) were removed, and proteins detected with two or more peptides were carried forward. Raw iBAQ intensities were normalized using median normalization. Median-normalized values were used for all analyses unless stated otherwise. All further data analysis was performed in the R statistical environment (v.4.2.1).

### EV isolation for biophysical studies

EVs were isolated from urines or cell line conditioned media for nanoparticle tracking analysis (NTA) and transmission electron microscopy (TEM) as described above, with the following changes. 5 mL of fluid was used for EV isolation using a SW55 Ti swinging bucket rotor (R_min_ 61, R_max_ 109, Beckman Coulter). To keep the k-factor consistent at each centrifugation step and taking into account the time needed for the rotor to achieve its desired speed (approximately 5 min), centrifugation time were reduced to 20 minutes and 1 hour for the 20,000 x *g* (k-factor 699) and 150,000 x *g* (k-factor 120) centrifugation steps, respectively. EV pellets were resuspended in 100-200 µL of cold, 0.22 µm filtered PBS, and stored at 4°C for no more than 16 hours prior to NTA or TEM analysis.

### Transmission electron microscopy

TEM was performed at SickKids Nanoscale Biomedical Imaging Facility and Princess Margaret Cancer Centre. Samples were deposited on formvar carbon-coated grids, washed once with water, and stained twice with uranyl acetate. Images were acquired on a Tecnai 20, Hitachi HT7800, and Talos™ F200X G2 transmission electron microscope. Images were processed with ImageJ (v.1.53t) for visualization.

### Nanoparticle tracking analysis

NTA for 34 men (**Table S1**) with uEV-P20 and uEV-P150 was performed using a NanoSight LM10 system configured with a 405 nm laser and a high sensitivity sCMOS camera. Camera settings were as follows: screen gain 3.0; camera level 11; 25 frames per second; slider gain 146. Each sample was diluted in particle-free PBS and introduced manually. Analysis was performed with NTA software (v.3.1 build 3.1.46). One technical replicate was captured per sample. For each replicate, three 30-second videos were captured, with approximately 20-200 particles in the field of view for each measurement. Ambient temperature was set at 22°C.

NTA for the DRE cohort (pre-DRE *vs.* post-DRE urine from two matched men) and cEVs (cEV-P20 and cEV-P150) was performed using a NanoSight NS300 system configured with a 405 nm laser and a high sensitivity sCMOS camera. Each sample was diluted in 0.22µm filtered PBS and introduced with a syringe pump at 60 μL/min. Analysis was performed with NTA software (v.3.4). For every sample, two to four technical replicates were captured. For each technical replicate, three 30 second videos were captured, with approximately 20-200 particles in the field of view for each measurement. Ambient temperature was set at 22°C.

Raw data files (“*filename-*ExperimentSummary.csv”) were parsed as follows for quantification and statistical analysis. Raw particle counts for each size bin were corrected for dilution factors, then grouped to biological replicates. Each data point represents the mean of all measurements for each biological replicate (**Table S1**). For cEVs, a processing replicate is defined as cEVs isolated from the same cell line at different passages. For uEVs, a biological replicate is defined as uEVs isolated from a specific biofluid (pre-DRE or post-DRE urine) from one individual. For visualizing particle concentration *vs.* size distribution for each replicate (**Figures S1D, 2C and S2D**) particle concentration was scaled with min-max [0,1] normalization with formula **Eq. 1**.

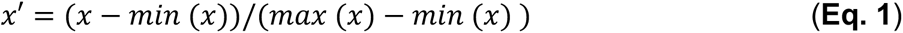

## QUANTIFICATION AND STATISTICAL ANALYSIS

Where appropriate, the quantitative analyses are described in the relevant sections of the Methods. Unless stated otherwise, bioinformatic and statistical analyses and plotting were performed using R (v.4.2.1). Data were visualized using R packages *BoutrosLab.plotting.general* (v.7.0.3)*, ggplot2* (v.3.4.0)*, ggbeeswarm* (v.0.6.0) *ggpubr* (v.0.4.0), and *ComplexHeatmap* (v.2.12.1). Qualitative variables were compared by Fisher Exact Test, and quantitative variables by Mann-Whitney U test for unpaired comparisons (*wilcox.test*), two-sided Wilcoxon signed-rank for pairwise comparisons (*wilcox.test*) and the Kruskal–Wallis test for multiple group comparisons. log_2_ fold change (log_2_FC) is calculated from the difference in medians. The specific statistical tests used are indicated in the figure legends. Multiple test *P-*values were adjusted using Benjamini-Hochberg method for independent tests unless stated otherwise. Correlation coefficients were determined by the Spearman method (*cor.test*). P-values for Spearman’s correlation are computed by asymptotic t approximation using an Edgeworth series. Statistical significance was set at P-value < 0.05. Missing values in protein-level data were replaced by random numbers drawn from the lower tail of the normal distribution 1.8 standard deviations from the mean (width = 0.2 standard deviations), unless stated otherwise^61^.

### Tissue specificity in DRE urines

To determine if DRE enriches for prostate tumor-derived proteins, proteomic profiles for the soluble protein (uSP) and uEV fractions were generated from urines collected pre- and post-DRE. Paired Student’s t-tests were used to identify differentially abundant proteins in pre- and post-DRE. To determine which proteins are anticipated to be derived from prostate tumors, proteins in this study were annotated based on detection in more than 10 tissue samples of two prostate cancer tissue proteomic datasets: Sinha *et. al.*^19^ (76 men with prostate cancer; 76 tumor samples) and Khoo *et. al.*^18^ (40 men with Prostate cancer; 81 samples [41 tumor and 40 NAT]); 7,438 proteins. To determine which proteins had elevated expression in human prostate tissue, the Human Protein Atlas (v.21.1, updated 2022-05-31) Human Tissue Specific Proteome^20^ from prostate was used to annotate proteins detected in this study. Proteins were included if they belonged to the ‘Tissue enriched’, ‘Group enriched’ or Tissue enhanced’ categories for the tissue of interest, totaling 127 genes for prostate tissue.

### Sample type correlations

Median protein abundances across samples were used for all comparisons using Spearman’s rank correlation (*cor.test*). P-values for Spearman’s correlation are computed by asymptotic t approximation using an Edgeworth series. Samples: uEV-P20 = 146, uEV-P150 = 148, Tissue = 157, uSP = 175. Proteins used for each comparison: uEV-P20 *vs.* uEV-P150 = 3,593; uSP *vs.* uEV-P20 = 2,839; uSP *vs.* uEV-P150 = 2,309; Tissue *vs.* uSP = 2,626; Tissue *vs.* uEV-P20 = 4,735; Tissue *vs.* uEV-P150 = 3,410.

### Identifying and annotating sample-type enriched proteins

To identify proteins enriched in uSP, uEV-P20 or uEV-P150 fractions, we considered both unique and differentially abundant proteins in each fraction. The set of fraction-unique proteins were defined as proteins detected in >90% of samples of one fraction and detected in less than 10% of samples of the other two fractions. Only samples with matched uSP, uEV-P20 and uEV-P150 fractions were used (288 samples from 96 patients). To identify differentially abundant proteins, proteins present in >20% of samples were used (Proteins: uSP = 2,909; uEV-P20 = 4,841, uEV-P150 = 3,389). A two-tailed, paired two-sided Wilcoxon signed-rank test was used to compare protein abundance in uSP *vs.* uEV-P20 (2,430 shared proteins), uSP *vs.* uEV-P150 (1,950 shared proteins), and uEV-P20 *vs.* uEV-P150 (2,939 shared proteins). Proteins were considered fraction-elevated if they were differentially abundant in both comparisons (**Figure 2I**). For example, proteins were considered uSP-unique if they were detected in >90% of uSP

samples, in <10% of uEV-P20 samples, and in <10% of uEV-P150 samples (**Figure 2H**). Proteins were considered ‘uSP-elevated’ if they were more abundant in uSP *vs.* uEV-P20 and uSP *vs.* uEV-P150 comparisons (FDR < 0.05 and |log_2_FC| > 0, intersect: 516 uSP-enriched proteins) (**Figure 2I**). This process resulted in a total of 606 uSP-enriched proteins (**Figure 2J**).

Fraction-enriched proteins were annotated with subcellular localization information from nine main categories (Secreted, Vesicles, Plasma membrane, Mitochondria, Cytosol, Nuclear Membrane, Nucleoplasm, Nucleoli and Golgi apparatus) from Human Protein Atlas’ Subcellular location data^55^ (v.22.0, proteinatlas.org). “Nuclear membrane”, “Nucleoplasm” and “Nucleoli” categories were collapsed into one category called “Nucleus”. Fisher’s Exact Test was used to test for over- or under-representation in each category for each fraction. The magnitude of the enrichment was estimated using the odds ratio (*epitools* v.0.5-10.1), with the union of proteins detected in fluids (uSP, uEV-P20 and uEV-P150; 6,540 proteins) used as a custom background.

### Sample-type tissue-enrichment scores

To score samples based on their tissue content, Gene Set Variation Analysis (v.1.44.5)^62^ was used to score samples using two custom gene signature sets – Prostate and Non-Prostate (Bladder + Kidney). For the signature gene sets, proteins that were enriched in each tissue type in the Genotype-Tissue Expression (GTEx) Project Bulk Tissue RNA-Seq dataset^25^ (V8; retrieved 2017-06-05; n_Prostate_ = 245 samples; n_Kidney_ = 89 samples; n_Bladder_ = 21 samples) and TCGA normal tissue adjacent to tumor from males^26–29^ (NAT; TCGA v.2016_01_28; n_PRAD_ = 52 samples, 19,821 genes; n_BLCA_ = 10 samples, 18,951 genes; n_KIRP_ = 22 samples, 19,518 genes; n_KIRC_ = 52 samples, 19,667 genes). For KIRP and KIRC NAT samples, we selected duplicated samples that had Spearman’s ρ > 0.99 and took the mean intensity, leaving a total of 71 KIRP/KIRC samples (19,829 genes). For each dataset, tissue-enriched genes were determined using Mann-Whitney U tests of each tissue of interest *vs.* the other two tissue types (*i.e.*, GTEx_Prostate_ *vs.* GTEx_Bladder + Kidney_; log_2_FC > 3 and FDR < 0.05). For the gene set signature, concordant genes in GTEx^25^ and TCGA NAT^26–29^ were selected (Genes: Prostate = 46, Bladder + Kidney = 41).

### Pathway analysis – sample type comparison

Proteins detected in >10 samples of each sample type (Proteins: Tissue = 7,438, uSP = 3,150, uEV-P20 = 5,462, uEV-P150 = 3,878) was used for pathway analysis using *gprofiler2* (v.0.2.1) against Gene Ontology:Cellular Component gene sets and visualized using EnrichmentMap (v.3.3.4) in Cytoscape (v.3.9.1).

### Cell line proteomics data

Cells and EVs were collected from three separate passages for each cell line, termed processing replicates, for proteomics. For each cell line and sample type, only proteins that were present in a minimum of two of three replicates were carried forward for analyses.

### Identifying temporally stable proteins

We generated proteomic profiles of uSP, uEV-P20 and uEV-P150 fractions from post-DRE urine from a longitudinal cohort composed of five patients. These patients had cISUP Grade Group 1 tumors and were on active surveillance (**Table S1**). None of the men upgraded in the time that the urines were collected. For statistical analysis of the longitudinal cohort, only reproducibly detected proteins were included for analysis. For each urine fraction, we selected proteins that were detected in at least two timepoints for each patient, and detected in at least two patients. This resulted in a total of 1,664 uSP proteins, 3,365 uEV-P20 proteins and 1,990 uEV-P150 proteins. The similarity of intra-patient and inter-patient proteomes were determined using Spearman’s correlation, calculated using the *cor()* function (*use* = “pairwise.complete.obs”) in *stats* package in R (v.4.2.1).

Identification of intra-variable and intra-stable proteins was performed as previously described^38^. Intra- and inter-individual variance in protein intensities was assessed using linear mixed-effects regression using the *lme4* package (v.1.1-31), and the intra-class correlation coefficient (ICC) was measured, which represents the proportion of inter-individual variance relative to total intra- and interindividual variance explained by a model. Proteins for which a model cannot be fitted due to random effect variances of close to zero. This resulted in a total of 1,664 uSP proteins, 3,365 uEV-P20 proteins and 1,990 uEV-P150 proteins with estimated ICC values.

### Prostate cancer vs NC comparison

Proteins detected in >50% of each sample type were considered for differential abundance analysis. This resulted in a total of 2,156 uSP proteins, 3,431 uEV-P20 proteins and 2,255 uEV-P150 proteins. Two-sided Mann-Whitney U test was used for comparisons.

### Feature selection and machine learning

To generate predictors that distinguishes men with prostate cancers and non-cancers (NC), proteomics data from uSP, uEV-P20, and uEV-P150 fractions were trained separately. Patients: uSP_NC_ = 39, uSP_PCa_= 136, uEV-P20_NC_ = 22, uEV-P20_PCa_ = 132, uEV-P150_NC_ = 25, uEV-P150_PCa_= 131. The same cohorts were used to generate predictors that distinguishes men with cISUP Grade Group (GG) 1 from men with cISUP Grade Group > 1 (uSP: 50 GG=1 *vs.* 61 GG>1; uEV-P20: 41 GG=1 *vs.* 63 GG>1; uEV-

P150: 40 GG=1 *vs.* 63 GG>1). To develop predictive models, all datasets were divided into two groups: feature selection (50% of dataset) and training (50% of dataset). Within each urinary fraction, proteins that were detected in more than 50% of all samples and temporally stable (intraclass coefficient > 0.4 from serial post-DRE urine collected from three cISUP GG1 patients at three timepoints) were passed into feature selection. Three methods were used to select the top 2 to 15 features within each dataset. For each feature, log_2_FC protein abundance was calculated and the significance level was assessed using the Mann-Whitney U test (*wilcox.test*). Features with the small*est P-val*ues were selected as the first set of top features. Features with the highest log_2_FC and P-value <0.001 were also selected as the second set of top features. Ten times repeated five-fold cross validated (*rfeControl*) was applied to get the third set of top features. Seven machine-learning algorithms were applied to the top features in the biomarker identification, including generalized linear models, random forest, k-nearest neighbour classification, naïve bayes, ridge, lasso, and elastic-net-regularized generalized linear model. Receiver operating characteristic (ROC) analysis with leave-one-out cross-validation was used to evaluate model performance with the use of ‘*pROC*’ package (v.1.18.0). Models with the highest area under the ROC (AUROC) were chosen and were fit to the entire dataset to get the final model. Machine learning algorithms were performed using the *caret* package (v.6.0.91) in R (v.3.6.1).

### Identifying prostate-specific, core and context-driven urine proteins

For each urine fraction (uEV-P20, uEV-P150 and uSP), we sought to identify proteins that were distinct to each urine fraction, prostate-derived, and indicative of disease state (prostate cancer *vs.* non-cancer or cISUP GG>1 *vs.* cISUP GG1). We also sought to identify fraction-specific, prostate-derived proteins that were stable in protein abundance, independent of disease state (*i.e.,* core proteome). Of the fraction-enriched proteins previously identified in **Figure 2J** (uEV-P20: 468 proteins, uEV-P150: 536 proteins, uSP: 606 proteins), we filtered for proteins that were detected in >50% of prostate tissue samples (*i.e.*, in more than 78 prostate tissues)^18,19^. We called these proteins fraction-enriched, prostate-derived proteins (uEV-P20: 431 proteins, uEV-P150: 437 proteins, uSP: 409 proteins). From this set of genes, we identified proteins that were stable across disease conditions, making up the core proteome – detected in >90% of samples in each fraction, not differentially abundant in prostate cancers *vs.* non-cancers (|log_2_FC| < 0.5), not differentially abundant in cISUP GG>1 *vs.* cISUP GG1 (|log_2_FC| < 0.5), and in the bottom 25% least variable proteins. This resulted in 64 uEV-P20 proteins, 107 uEV-P150 proteins, and 78 uSP proteins. We also identified proteins that were differentially abundant in prostate cancers *vs.* non-cancers (*i.e.*, “Tumor markers”; |log_2_FC| > 1 and FDR < 0.05) and in cISUP Grade Group >1 *vs.* cISUP Grade Group 1 (*i.e.*, “Grade markers”; |log_2_FC| > 1 and unadjusted p-value < 0.05). From each of these groups – Core, Tumor, and Grade markers – we identified proteins with predicted cell surface localization (Surface Prediction Consensus [SPC] score^41^ > 2) that could serve as potential markers for each of these groups.

## ACKNOWLEDGEMENTS

We thank the Nanoscale Biomedical Imaging Facility at SickKids and Mohammad Mazhab-Jafari at Princess Margaret Cancer Centre for acquiring the negative stain transmission electron microscopy images. We would also like to thank the SickKids Structural & Biophysical Core Facility for allowing use of their instruments for nanoparticle tracking analysis. The Genotype-Tissue Expression (GTEx) Project was supported by the Common Fund of the Office of the Director of the National Institutes of Health, and by NCI, NHGRI, NHLBI, NIDA, NIMH, and NINDS. The results here are in whole or part based upon data generated by the TCGA Research Network: https://www.cancer.gov/tcga. A.K., M.G., and L.Y.L are supported by the Ontario Graduate Scholarship and Ontario Student Opportunity Trust Fund Awards. L.Y.L is supported by a Vanier Canada Graduate Scholarship. This study was supported by the National Institutes of Health through awards U01CA214194, U2CCA271894, U24CA248265, R01CA272678, P20CA252717 and UM1TR004360. This work was supported by a Prostate Cancer Foundation Special Challenge Award to P.C.B. (20CHAS01) made possible by the generosity of Mr. Larry Ruvo. This study was supported by Canadian Institutes of Health Research Project Grants to T.K. (PJT156357) and S.K.L. (PJT162384). T.K. was supported through the Canadian Research Chair program. T.K., S.K.L. and D.V. also received support through a Prostate Cancer Canada Discovery Grant (D2019-2113).

## AUTHOR CONTRIBUTIONS

A.K., M.G., A.M., Al.K, B.P.M., R.S.L, L.Y., M.D., D.V., S.K.L., O.J.S. and J.O.N. contributed to data acquisition. A.K. and A.M. acquired the mass spectrometry data. A.K., Z.Q., M.G., L.Y.L and V.I. analyzed the data. S.K.L., O.J.S., J.O.N., P.C.B., and T.K. supervised the study. A.K., M.G., Z.Q., M.W., L.Y.L., T.K. and P.C.B wrote the manuscript, which all other authors edited and approved.

## RESOURCE AVAILABILITY

### Lead contact

Further information and requests for resources and reagents should be directed to and will be fulfilled by the lead contact, Thomas Kislinger.

### Materials availability

This study did not generate new unique reagents.

### Data and code availability

- All mass spectrometry raw files and processed result files acquired in this study are publicly available from UCSD’s MassIVE database (ftp://massive.ucsd.edu) under dataset identifier #MSV000092061. Processed proteomics data are available in this paper’s **Table S2**.
- We have submitted all relevant data of our EV experiments to the EV-TRACK knowledgebase^52^ (EV-TRACK ID: EV230578).
- A web application (http://kislingerlab.uhnres.utoronto.ca/ev/home/) has also been created for browsing data.
- Any additional information required to reanalyze the data reported in this paper is available from the lead contact upon request.

## ADDITIONAL RESOURCES

An accompanying website for this study provides an interactive browser for interrogation of multiple datasets and clinical cohorts, available at http://kislingerlab.uhnres.utoronto.ca/ev/home/.

## EXPERIMENTAL MODEL AND SUBJECT DETAILS

### Human subjects

Samples were obtained from men following informed consent and use of Institutional Review Board approved protocols at Eastern Virginia Medical School (EVMS, Norfolk, Virginia, USA, IRB# 06-12-FB-0343), Sunnybrook Health Sciences Centre (SHSC, Toronto, Ontario, Canada, Project #2457) and the Research Ethics Review Board at the University Health Network (UHN, Toronto, Ontario, Canada, 10-0159 and 19-5009). Men with benign prostatic conditions (**Table S1**) included individuals with elevated serum PSA (sPSA) levels but no diagnosed prostate cancer on transrectal ultrasound-guided 12-core biopsy (Biopsy-negative; 20 patients; median sPSA 5.2 ng/mL, range 0.5 – 31.5 ng/mL), or benign prostatic hyperplasia (BPH; 44 patients; median sPSA 6.3 ng/mL, range 1.7 – 11.9 ng/mL). Selection criteria for men with benign prostatic conditions included a diagnostic sPSA level < 20 ng/mL and post-surgery sPSA level <0.1 ng/mL to exclude highly metastatic men. Other clinical details are detailed in **Table S1**.

### Urine collection

The first 15 mL of first-catch urine collected post digital rectal exam (DRE) (post-DRE urine) was collected by performing a gentle massage of the prostate gland during DRE prior to biopsy, as previously described^22^. For the DRE cohort (**Table S1**), which comprised of ten men with clinical ISUP Grade Group (cISUP GG) 1 tumors, mid-stream urine was collected an hour before the DRE massage (pre-DRE urine). Matched post-DRE urine was also collected for these ten men. The longitudinal cohort comprised of five men with cISUP Grade Group 1 tumors who are on active surveillance and did not upgrade in the period of 12-16 months after their first DRE (**Table S1**). Serial post-DRE urine was collected for each patient at three timepoints. Each timepoint was 3-12 months apart. Pre- and post-DRE urine was centrifuged at 2,000 x *g* for 15 min at 4°C to pellet cellular debris, and the resulting urine supernatant was stored at −80°C.

### Cell lines

Commercial human prostate cell lines DU145, PC3, 22Rv1, LNCaP and RWPE1 were acquired from ATCC. All cell lines are immortalized cell lines from males. Cell line identity was confirmed by short tandem repeat testing. Mycoplasma negativity was confirmed using the Universal Mycoplasma Testing kit (ATCC). Cells were seeded in two T500 Nunc^TM^ TripleFlasks^TM^ (total area = 1,000 cm^2^) with 100 mL of media or ten 15 cm plates (total area = 1,480 cm^2^) in 20 mL of media each and cultured in a 37°C incubator with 5% CO_2_. RPMI media (Gibco) supplemented with 10% fetal bovine serum and 1% penicillin-streptomycin-glutamine (PSG) was used for the prostate cancer cell lines (DU145, PC3, 22Rv1 and LNCaP) and Keratinocyte-serum free media supplemented with 0.05 mg/mL BPE, 5 ng/mL EGF, and 1% PSG was used for the RWPE1 cell line.

## SUPPLEMENTARY TABLES

**Table S1.Patient characteristics, related to all Figures and Methods.**

**Table S2. Processed proteomic profiles of urines, uEVs and cEVs, related to all Figures and Methods.**

**Table S3. Results from pathway analysis, related to Figures 2K, and 3E.**

**Table S4. Results from statistical analyses, related to all Figures and Methods.**

## FIGURE LEGENDS

**Figure S1.**
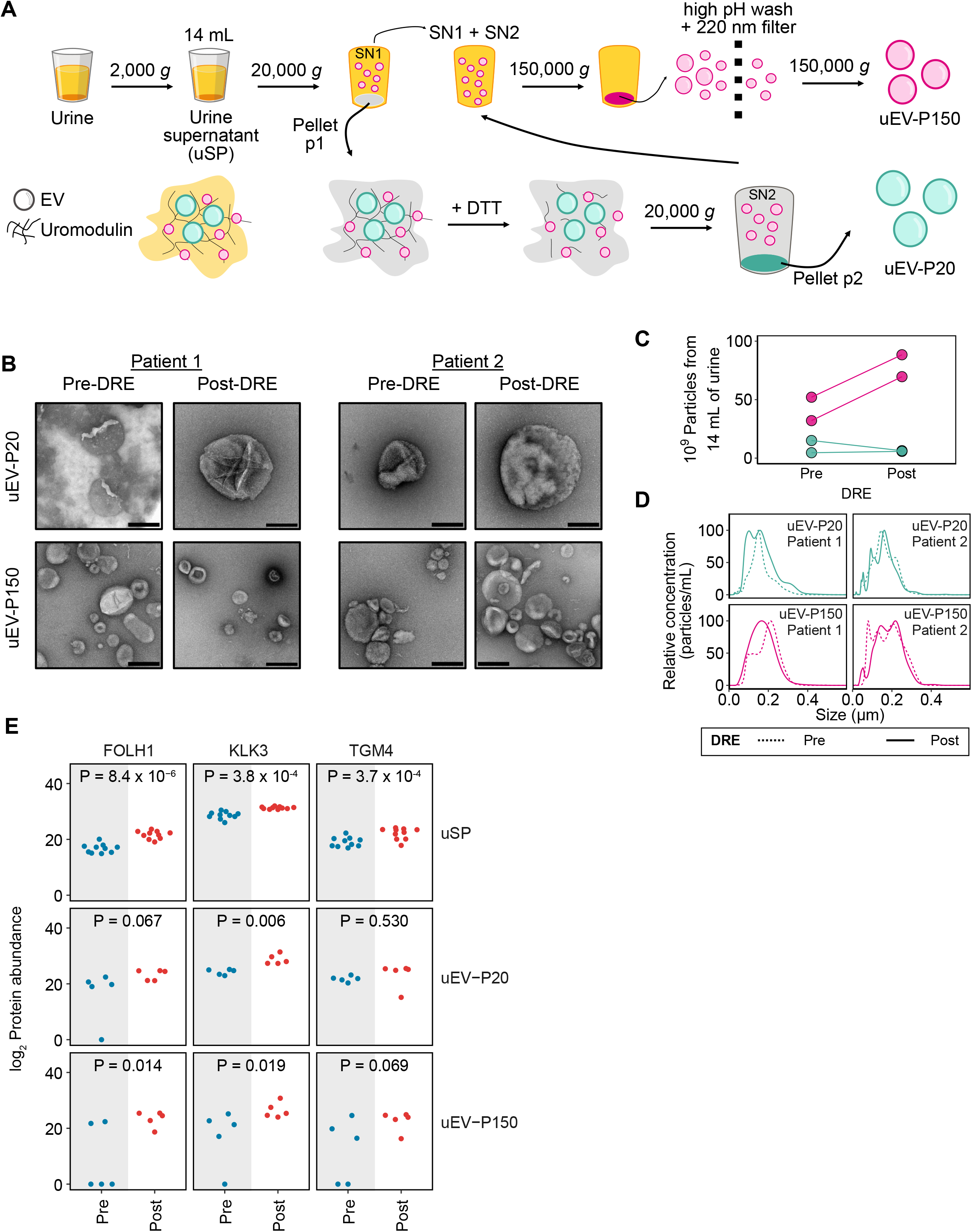
Molecular and biophysical characterization of uEVs, related to Figure 1. **(A)** Urinary extracellular vesicle isolation by differential ultracentrifugation and filtration. **(B)** Negative stain TEM images of uEVs isolated from pre- and post-DRE urine from two men. Scale bar: 200 nm. **(C)** Number of uEVs isolated from 14 mL of urine from two men with cISUP GG 1 tumors. **(D)** Size distribution of uEV-P20 and uEV-P150 from two matched patients by nanoparticle tracking analysis. **(E)** Protein abundance for prostate marker proteins in pre- *vs.* post-DRE urine. P values from a Wilcoxon signed-rank test.

**Figure S2.**
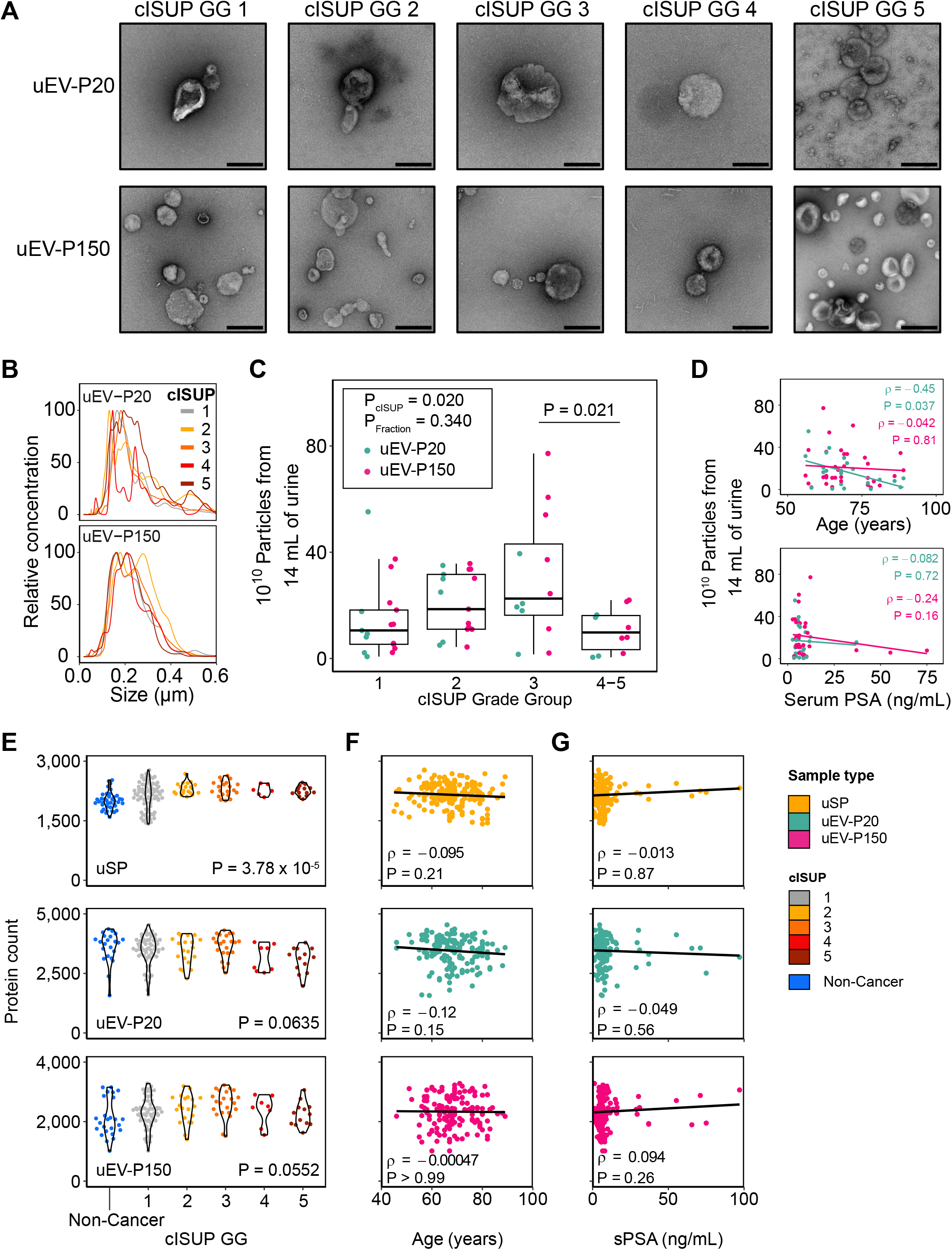
uEV associations with clinical covariates, related to Figure 2. **(A)** Negative stain TEM images of uEVs isolated from the post-DRE urine of men with different cISUP Grade Group (GG) tumors. Scale bar: 200 nm. **(B)** Size distribution by NTA. Each curve represents the mean size for each cISUP GG. **(C)** Number of uEV-P20 and uEV-P150 isolated from 14 mL of post-DRE urine from men with different cISUP Grade Group tumors. Patients: uEV-P20_GG1_ = 7, uEV-P150_GG1_ = 10, uEV-P20_GG2_ = 6, uEV-P20_GG2_ = 11, uEV-P20_GG3_ = 5, uEV-P150_GG3_ = 7, uEV-P20_GG4-5_ = 4, uEV-P150_GG4-5_ = 6. Significantly different groups (two-way ANOVA with *post hoc* Tukey’s HSD < 0.05) are labeled. **(D)** Number of uEV particles and age at diagnosis (top) or serum PSA level (bottom). **(E-G)** Associations between protein counts and **(E)** cISUP GG, **(F)** age and **(G)** serum PSA. P-values in (E) from one-way ANOVA. Spearman’s ρ for (F-G).

**Figure S3.**
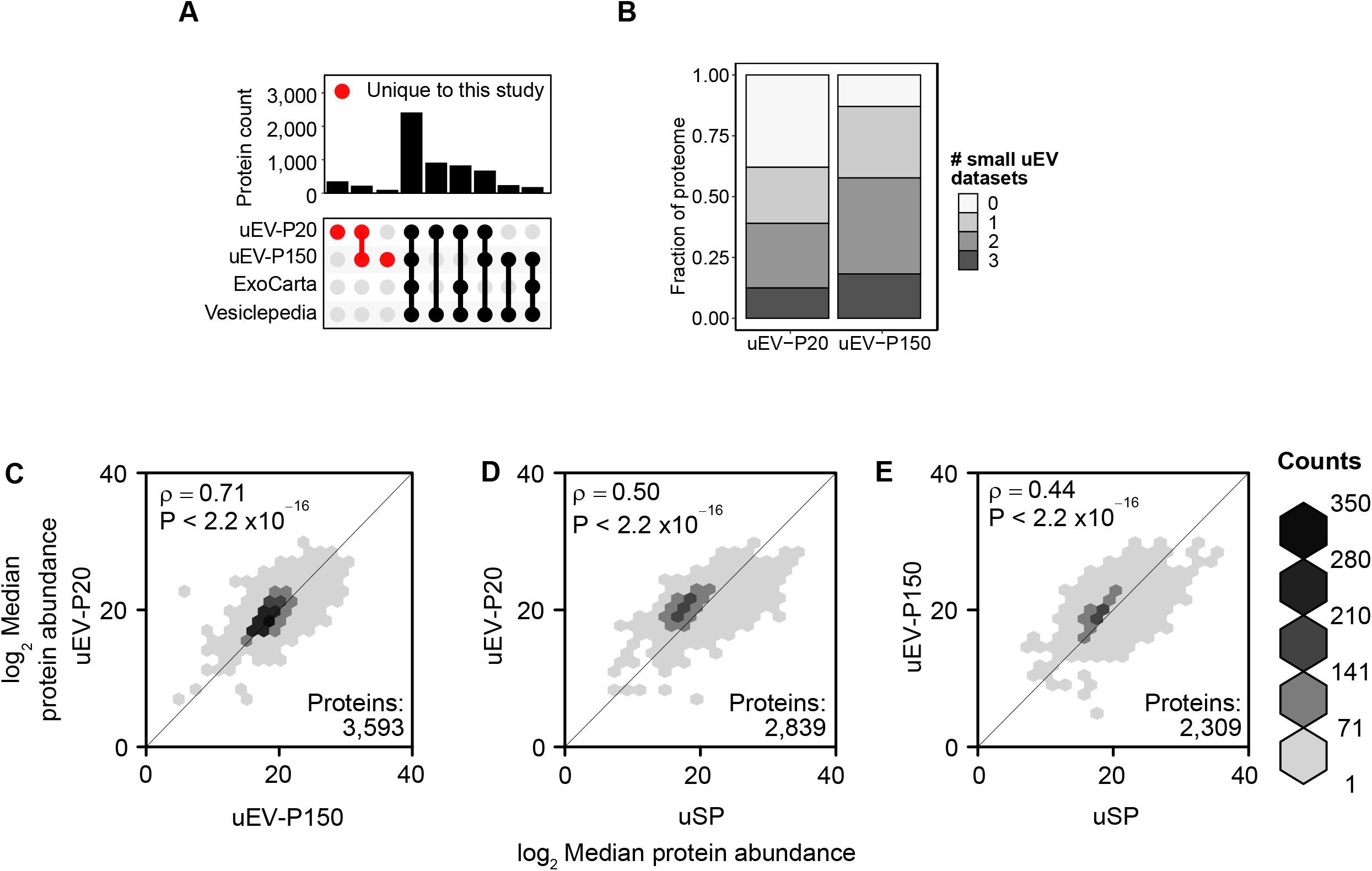
uEV proteomes are distinct from uSP proteomes, related to Figure 2. **(A)** Overlap in uEV proteins detected in this study with ExoCarta^23^ and Vesiclepedia^24^. Proteins unique to the current study are in red. **(B)** Fraction of uEV proteins detected in this dataset that were also detected in three other published post-DRE urine small uEV datasets^17,21,22^. **(C-E)** Median protein abundance showing sample type correlation (Spearman’s ρ) between **(C)** uEV-P20 *vs.* uEV-P150, **(D)** uEV-P20 *vs.* uSP and **(E)** uEV-P150 *vs.* uSP.

**Figure S4.**
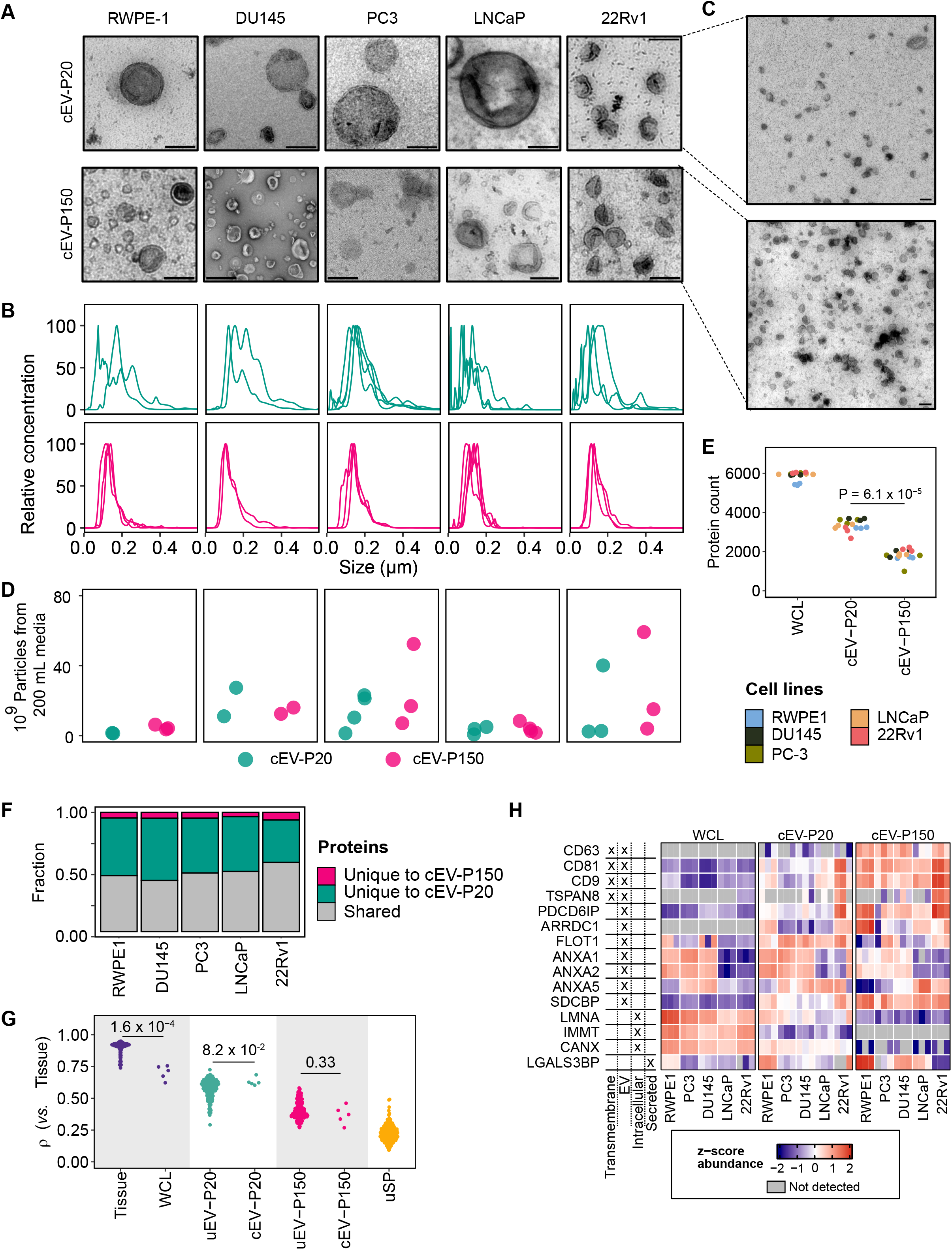
Biophysics and proteomics of cell line-derived EVs, related to Figure 4. (A) Negative stain TEM images of cell line EVs (cEV) isolated from conditioned media of prostate normal epithelium cell line (RWPE-1) and prostate cancer cell lines (DU145, PC3, LNCaP and 22Rv1). Scale bar: 200 nm. (B) Size distribution determined by nanoparticle tracking analysis. Each curve represents the mean size for each biological replicate of cEV-P20 (top) or cEV-P150 (bottom). (C) Wide-view negative stain TEM images of cEV-P20 (top) and cEV-P150 (bottom) isolated from conditioned media of 22Rv1 prostate cancer cell line. Scale bar: 200 nm. (D) Number of cEV-P20 and cEV-P150 particles determined by nanoparticle tracking analysis. Replicates: cEV-P20_RWPE1_ = 2, cEV-P150_RWPE1_ = 3; cEV-P20_DU145_ = 2, cEV-P150_DU145_ = 2; cEV-P20_PC3_ = 4, cEV-P150_PC3_ = 3; cEV-P20_LNCaP_ = 3, cEV-P150_LNCaP_ = 4; cEV-P20_22Rv1_ = 3, cEV-P150_22Rv1_ = 3. (E) Protein counts for each cell line-derived sample type, colored by cell line. Processing triplicates are shown. P-values from two-tailed paired Wilcoxon test. (F) Fraction of shared or unique proteins between cEV-P20 and cEV-P150 fractions across cell lines. (G) Spearman’s ρ between log_2_ mean protein abundance in prostate tissue (157 samples), and individual samples from tissue, whole cell lysate (WCL, 5 cell lines), uEVs (n_uEV-P20_ = 146 samples, n_cEV-P20_ = 5 cell lines, n_uEV-P150_ = 148, n_cEV-P150_ = 5 cell lines), and soluble proteins (uSP, n = 175). Each data point from cell line samples represents the mean of 3 processing replicates per cell line. (H) Z-scores of log_2_ protein abundance of select EV and non-EV markers (MISEV 2018^63^) in WCL, cEV-P20 and cEV-P150.

**Figure S5.**
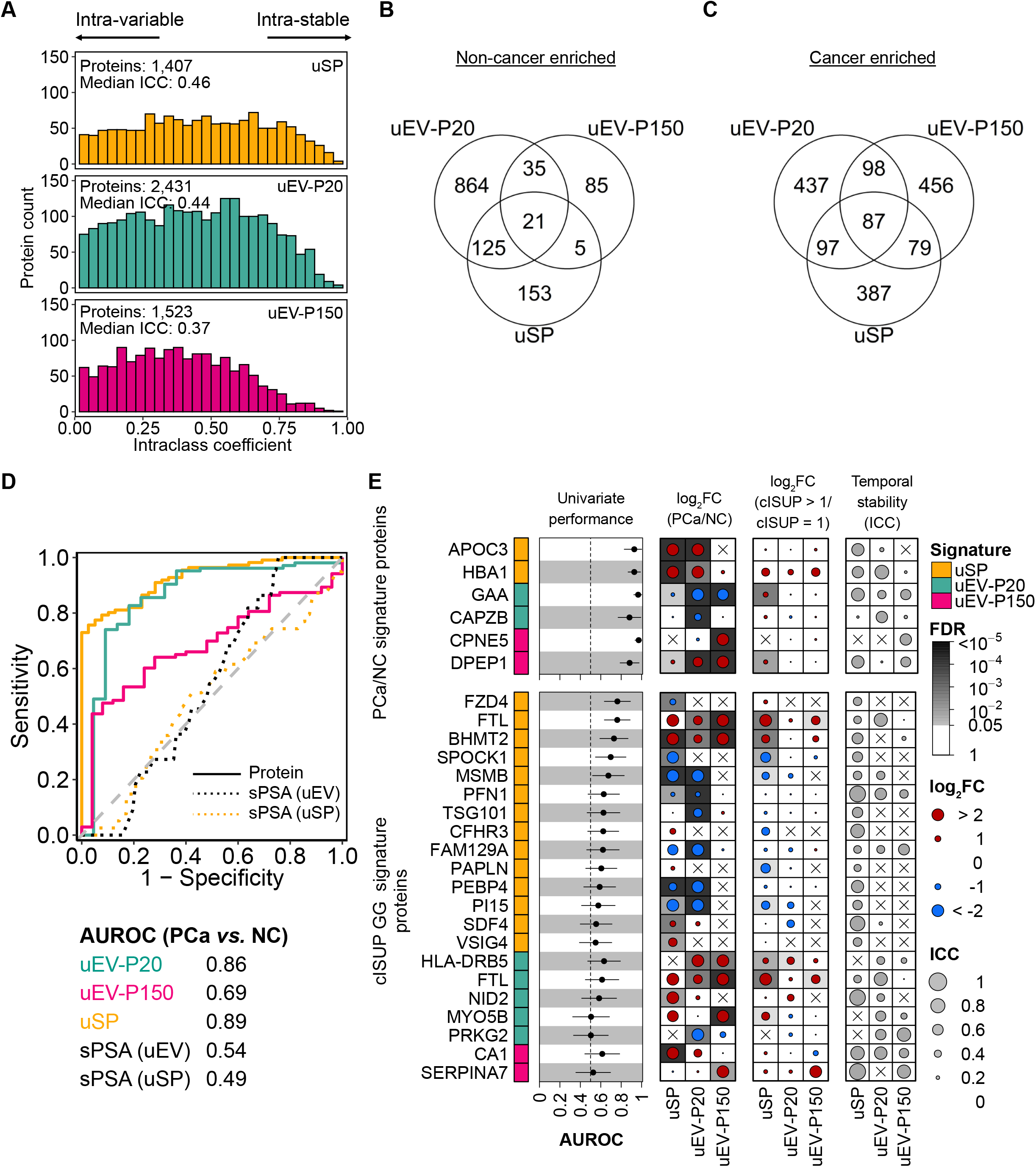
Prostate cancer *vs.* non-cancer proteomes across sample types, related to Figure 5. **(A)** Distribution of estimated variability for proteins in each urine fraction, estimated using intraclass correlation coefficient. **(B-C)** Overlap of proteins significantly upregulated in **(B)** non-cancer fractions and **(C)** prostate cancer fractions using a Mann-Whitney U test FDR < 0.05. **(D)** ROC curve of cISUP GG protein model performance in classifying prostate cancers (PCa) *vs.* non-cancers (NC). Dotted lines show performance of serum PSA for uEVs (patients with matched uEV-P20 and uEV-P150) and uSP. **(E)** The performance (AUROC) for all signature proteins in a univariate model (left). Lines denote 95% CI. Dotmap summarizing differences in protein abundance (log_2_FC) in patients with prostate cancer and non-cancers, and patients with cISUP GG>1 or GG1 tumors. Dotmap on right gives intraclass coefficient (right) for each of these proteins from a cISUP GG1 patient cohort (5 active surveillance patients, no upgrading, 3 timepoints over 12-24 months). Grey background in dotmap denotes FDR < 0.05. Crosses denote proteins not detected in at least half of samples in that urine fraction.

**Figure S6.**
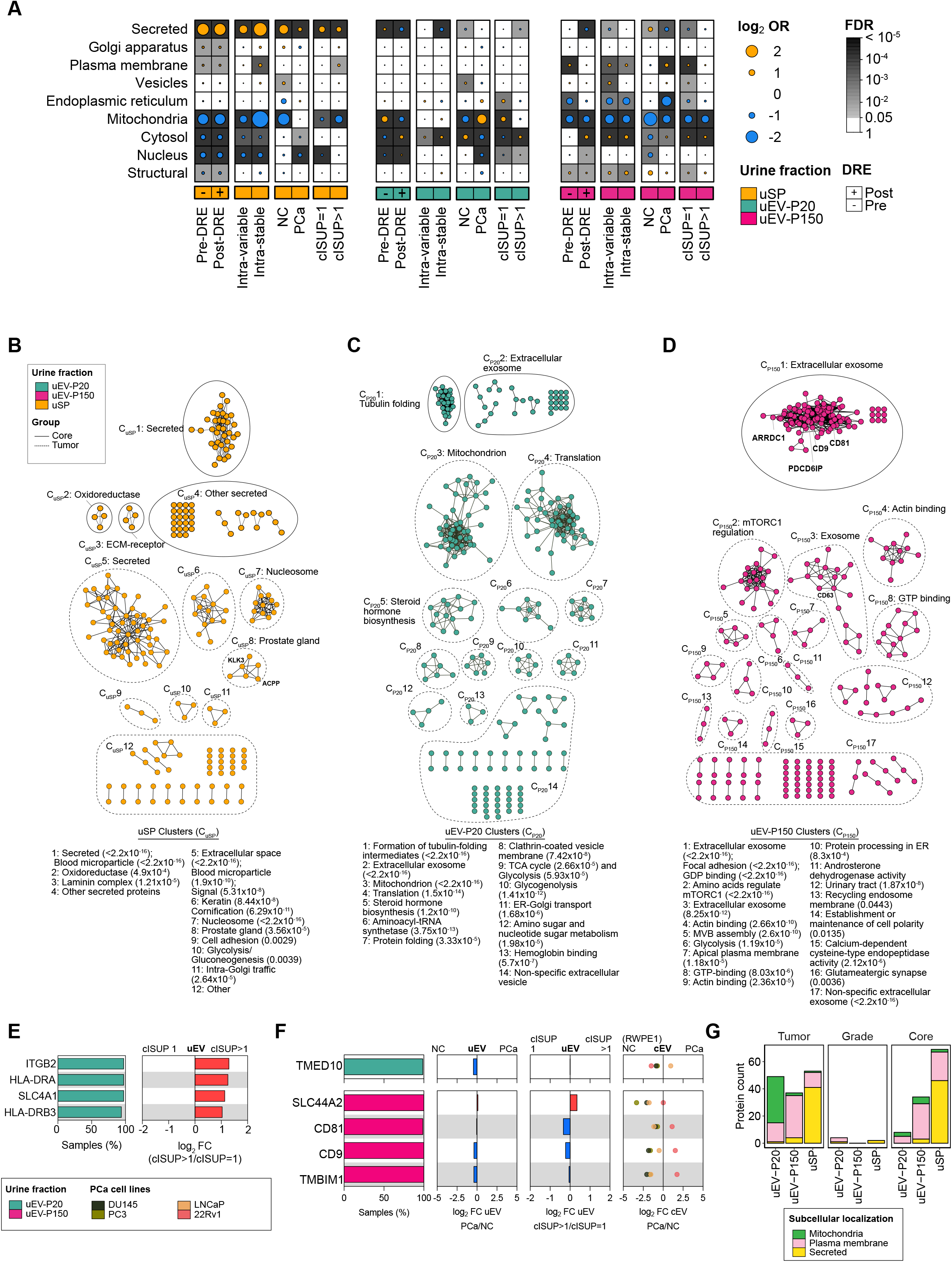
Pathways implicated in EV cargo dysregulation, related to Figure 6. **(A)** Dot plot of log_2_ odds ratio of over-representation for each term. Background shading indicates significant terms (FDR < 0.05, Fisher’s Exact test). **(B-D)** STRING and pathway analysis of proteins in each fraction belonging to core or tumor gene sets (*i.e.*, differentially abundant in PCa/NC). Nodes represent proteins, and edges represent protein-protein associations^64^. Each cluster is annotated with pathway terms and P-value in parentheses. **(E)** Summary of uEV-P20 grade markers with predicted cell surface localization from Figure 6C. Proteins are annotated with the frequency of detection in each fraction (left panel), and differential abundance in cISUP GG>1 *vs.* cISUP GG1 (right panel). **(F)** Summary of core uEV proteins with predicted cell surface localization from Figure 6C, annotated with the frequency of detection in each fraction (left panel), differential abundance in PCa *vs.* NC uEVs, cISUP GG uEVs, and PCa *vs.* NC cEVs. uEV: Urinary EV, cEV: cell line EV. **(G)** Subcellular annotation^55^ of all disease-specific and core markers for all fractions.

## REFERENCES

1. Ridley, J.W. (2018). Fundamentals of the study of urine and body fluids (Springer).

2. Drake, R.R., White, K.Y., Fuller, T.W., Igwe, E., Clements, M.A., Nyalwidhe, J.O., Given, R.W., Lance, R.S., and Semmes, O.J. (2009). Clinical collection and protein properties of expressed prostatic secretions as a source for biomarkers of prostatic disease. J. Proteomics 72, 907–917. 10.1016/j.jprot.2009.01.007.

3. Simerville, J.A., Maxted, W.C., and Pahira, J.J. (2005). Urinalysis: a comprehensive review. Am. Fam. Physician 71, 1153–1162.

4. Nagaraj, N., and Mann, M. (2011). Quantitative analysis of the intra- and inter-individual variability of the normal urinary proteome. J. Proteome Res. 10, 637–645. 10.1021/pr100835s.

5. Tomlins, S.A., Day, J.R., Lonigro, R.J., Hovelson, D.H., Siddiqui, J., Kunju, L.P., Dunn, R.L., Meyer, S., Hodge, P., Groskopf, J., et al. (2016). Urine TMPRSS2:ERG Plus PCA3 for Individualized Prostate Cancer Risk Assessment. Eur. Urol. 70, 45–53. 10.1016/j.eururo.2015.04.039.

6. Khoo, A., Liu, L.Y., Nyalwidhe, J.O., Semmes, O.J., Vesprini, D., Downes, M.R., Boutros, P.C., Liu, S.K., and Kislinger, T. (2021). Proteomic discovery of non-invasive biomarkers of localized prostate cancer using mass spectrometry. Nat. Rev. Urol. 18, 707–724. 10.1038/s41585-021-00500-1.

7. McKiernan, J., Donovan, M.J., Margolis, E., Partin, A., Carter, B., Brown, G., Torkler, P., Noerholm, M., Skog, J., Shore, N., et al. (2018). A Prospective Adaptive Utility Trial to Validate Performance of a Novel Urine Exosome Gene Expression Assay to Predict High-grade Prostate Cancer in Patients with Prostate-specific Antigen 2-10ng/ml at Initial Biopsy. Eur. Urol. 74, 731–738. 10.1016/j.eururo.2018.08.019.

8. Tosoian, J.J., Trock, B.J., Morgan, T.M., Salami, S.S., Tomlins, S.A., Spratt, D.E., Siddiqui, J., Kunju, L.P., Botbyl, R., Chopra, Z., et al. (2021). Use of the MyProstateScore Test to Rule Out Clinically Significant Cancer: Validation of a Straightforward Clinical Testing Approach. J. Urol. 205, 732–739. 10.1097/JU.0000000000001430.

9. Mishra, J., Dent, C., Tarabishi, R., Mitsnefes, M.M., Ma, Q., Kelly, C., Ruff, S.M., Zahedi, K., Shao, M., Bean, J., et al. (2005). Neutrophil gelatinase-associated lipocalin (NGAL) as a biomarker for acute renal injury after cardiac surgery. Lancet Lond. Engl. 365, 1231–1238. 10.1016/S0140-6736(05)74811-X.

10. Soloway, M.S., Briggman, V., Carpinito, G.A., Chodak, G.W., Church, P.A., Lamm, D.L., Lange, P., Messing, E., Pasciak, R.M., Reservitz, G.B., et al. (1996). Use of a new tumor marker, urinary NMP22, in the detection of occult or rapidly recurring transitional cell carcinoma of the urinary tract following surgical treatment. J. Urol. 156, 363–367. 10.1097/00005392-199608000-00008.

11. Decramer, S., Gonzalez de Peredo, A., Breuil, B., Mischak, H., Monsarrat, B., Bascands, J.-L., and Schanstra, J.P. (2008). Urine in clinical proteomics. Mol. Cell. Proteomics MCP 7, 1850–1862. 10.1074/mcp.R800001-MCP200.

12. Julian, B.A., Suzuki, H., Suzuki, Y., Tomino, Y., Spasovski, G., and Novak, J. (2009). Sources of Urinary Proteins and their Analysis by Urinary Proteomics for the Detection of Biomarkers of Disease. Proteomics Clin. Appl. 3, 1029–1043. 10.1002/prca.200800243.

13. van Niel, G., D’Angelo, G., and Raposo, G. (2018). Shedding light on the cell biology of extracellular vesicles. Nat. Rev. Mol. Cell Biol. 19, 213–228. 10.1038/nrm.2017.125.

14. Xu, R., Rai, A., Chen, M., Suwakulsiri, W., Greening, D.W., and Simpson, R.J. (2018). Extracellular vesicles in cancer - implications for future improvements in cancer care. Nat. Rev. Clin. Oncol. 15, 617–638. 10.1038/s41571-018-0036-9.

15. Kalluri, R., and LeBleu, V.S. (2020). The biology, function, and biomedical applications of exosomes. Science 367, eaau6977. 10.1126/science.aau6977.

16. Pellegrini, K.L., Patil, D., Douglas, K.J.S., Lee, G., Wehrmeyer, K., Torlak, M., Clark, J., Cooper, C.S., Moreno, C.S., and Sanda, M.G. (2017). Detection of prostate cancer-specific transcripts in extracellular vesicles isolated from post-DRE urine. The Prostate 77, 990–999. 10.1002/pros.23355.

17. Correll, V.L., Otto, J.J., Risi, C.M., Main, B.P., Boutros, P.C., Kislinger, T., Galkin, V.E., Nyalwidhe, J.O., Semmes, O.J., and Yang, L. (2022). Optimization of small extracellular vesicle isolation from expressed prostatic secretions in urine for in-depth proteomic analysis. J. Extracell. Vesicles 11, e12184. 10.1002/jev2.12184.

18. Khoo, A., Liu, L.Y., Sadun, T.Y., Salmasi, A., Pooli, A., Felker, E., Houlahan, K.E., Ignatchenko, V., Raman, S.S., Sisk, A.E., et al. (2022). Prostate cancer multiparametric magnetic resonance imaging visibility is a tumor-intrinsic phenomena. J. Hematol. Oncol.J Hematol Oncol 15, 48. 10.1186/s13045-022-01268-6.

19. Sinha, A., Huang, V., Livingstone, J., Wang, J., Fox, N.S., Kurganovs, N., Ignatchenko, V., Fritsch, K., Donmez, N., Heisler, L.E., et al. (2019). The Proteogenomic Landscape of Curable Prostate Cancer. Cancer Cell 35, 414–427.e6. 10.1016/j.ccell.2019.02.005.

20. Uhlén, M., Fagerberg, L., Hallström, B.M., Lindskog, C., Oksvold, P., Mardinoglu, A., Sivertsson, Å., Kampf, C., Sjöstedt, E., Asplund, A., et al. (2015). Proteomics. Tissue-based map of the human proteome. Science 347, 1260419. 10.1126/science.1260419.

21. Dhondt, B., Geeurickx, E., Tulkens, J., Van Deun, J., Vergauwen, G., Lippens, L., Miinalainen, I., Rappu, P., Heino, J., Ost, P., et al. (2020). Unravelling the proteomic landscape of extracellular vesicles in prostate cancer by density-based fractionation of urine. J Extracell Vesicles 9, 1736935. 10.1080/20013078.2020.1736935.

22. Principe, S., Jones, E.E., Kim, Y., Sinha, A., Nyalwidhe, J.O., Brooks, J., Semmes, O.J., Troyer, D.A., Lance, R.S., Kislinger, T., et al. (2013). In-depth proteomic analyses of exosomes isolated from expressed prostatic secretions in urine. Proteomics 13, 1667–1671. 10.1002/pmic.201200561.

23. Keerthikumar, S., Chisanga, D., Ariyaratne, D., Al Saffar, H., Anand, S., Zhao, K., Samuel, M., Pathan, M., Jois, M., Chilamkurti, N., et al. (2016). ExoCarta: A Web-Based Compendium of Exosomal Cargo. J. Mol. Biol. 428, 688–692. 10.1016/j.jmb.2015.09.019.

24. Kalra, H., Simpson, R.J., Ji, H., Aikawa, E., Altevogt, P., Askenase, P., Bond, V.C., Borràs, F.E., Breakefield, X., Budnik, V., et al. (2012). Vesiclepedia: a compendium for extracellular vesicles with continuous community annotation. PLoS Biol. 10, e1001450. 10.1371/journal.pbio.1001450.

25. GTEx Consortium (2020). The GTEx Consortium atlas of genetic regulatory effects across human tissues. Science 369, 1318–1330. 10.1126/science.aaz1776.

26. Cancer Genome Atlas Research Network (2015). The Molecular Taxonomy of Primary Prostate Cancer. Cell 163, 1011–1025. 10.1016/j.cell.2015.10.025.

27. Cancer Genome Atlas Research Network (2014). Comprehensive molecular characterization of urothelial bladder carcinoma. Nature 507, 315–322. 10.1038/nature12965.

28. Cancer Genome Atlas Research Network, Linehan, W.M., Spellman, P.T., Ricketts, C.J., Creighton, C.J., Fei, S.S., Davis, C., Wheeler, D.A., Murray, B.A., Schmidt, L., et al. (2016). Comprehensive Molecular Characterization of Papillary Renal-Cell Carcinoma. N. Engl. J. Med. 374, 135–145. 10.1056/NEJMoa1505917.

29. Cancer Genome Atlas Research Network (2013). Comprehensive molecular characterization of clear cell renal cell carcinoma. Nature 499, 43–49. 10.1038/nature12222.

30. Tien, W.-S., Chen, J.-H., and Wu, K.-P. (2017). SheddomeDB: the ectodomain shedding database for membrane-bound shed markers. BMC Bioinformatics 18, 42. 10.1186/s12859-017-1465-7.

31. Adachi, J., Kumar, C., Zhang, Y., Olsen, J.V., and Mann, M. (2006). The human urinary proteome contains more than 1500 proteins, including a large proportion of membrane proteins. Genome Biol. 7, R80. 10.1186/gb-2006-7-9-R80.

32. Tricarico, C., Clancy, J., and D’Souza-Schorey, C. (2017). Biology and biogenesis of shed microvesicles. Small GTPases 8, 220–232. 10.1080/21541248.2016.1215283.

33. Lischnig, A., Bergqvist, M., Ochiya, T., and Lässer, C. (2022). Quantitative Proteomics Identifies Proteins Enriched in Large and Small Extracellular Vesicles. Mol. Cell. Proteomics MCP 21, 100273. 10.1016/j.mcpro.2022.100273.

34. Lázaro-Ibáñez, E., Lunavat, T.R., Jang, S.C., Escobedo-Lucea, C., Oliver-De La Cruz, J., Siljander, P., Lötvall, J., and Yliperttula, M. (2017). Distinct prostate cancer-related mRNA cargo in extracellular vesicle subsets from prostate cell lines. BMC Cancer 17, 92. 10.1186/s12885-017-3087-x.

35. Sardana, G., Jung, K., Stephan, C., and Diamandis, E.P. (2008). Proteomic analysis of conditioned media from the PC3, LNCaP, and 22Rv1 prostate cancer cell lines: discovery and validation of candidate prostate cancer biomarkers. J. Proteome Res. 7, 3329–3338. 10.1021/pr8003216.

36. Escola, J.M., Kleijmeer, M.J., Stoorvogel, W., Griffith, J.M., Yoshie, O., and Geuze, H.J. (1998). Selective enrichment of tetraspan proteins on the internal vesicles of multivesicular endosomes and on exosomes secreted by human B-lymphocytes. J. Biol. Chem. 273, 20121–20127. 10.1074/jbc.273.32.20121.

37. Théry, C., Regnault, A., Garin, J., Wolfers, J., Zitvogel, L., Ricciardi-Castagnoli, P., Raposo, G., and Amigorena, S. (1999). Molecular characterization of dendritic cell-derived exosomes. Selective accumulation of the heat shock protein hsc 73. J. Cell Biol. 147, 599–610. 10.1083/jcb.147.3.599.

38. Jeon, J., Olkhov-Mitsel, E., Xie, H., Yao, C.Q., Zhao, F., Jahangiri, S., Cuizon, C., Scarcello, S., Jeyapala, R., Watson, J.D., et al. (2020). Temporal Stability and Prognostic Biomarker Potential of the Prostate Cancer Urine miRNA Transcriptome. J. Natl. Cancer Inst. 112, 247–255. 10.1093/jnci/djz112.

39. Fraser, M., Sabelnykova, V.Y., Yamaguchi, T.N., Heisler, L.E., Livingstone, J., Huang, V., Shiah, Y.-J., Yousif, F., Lin, X., Masella, A.P., et al. (2017). Genomic hallmarks of localized, non-indolent prostate cancer. Nature 541, 359–364. 10.1038/nature20788.

40. Hoshino, A., Kim, H.S., Bojmar, L., Gyan, K.E., Cioffi, M., Hernandez, J., Zambirinis, C.P., Rodrigues, G., Molina, H., Heissel, S., et al. (2020). Extracellular Vesicle and Particle Biomarkers Define Multiple Human Cancers. Cell 182, 1044–1061.e18. 10.1016/j.cell.2020.07.009.

41. Waas, M., Snarrenberg, S.T., Littrell, J., Jones Lipinski, R.A., Hansen, P.A., Corbett, J.A., and Gundry, R.L. (2020). SurfaceGenie: a web-based application for prioritizing cell-type-specific marker candidates. Bioinformatics 36, 3447–3456. 10.1093/bioinformatics/btaa092.

42. Dixson, A.C., Dawson, T.R., Di Vizio, D., and Weaver, A.M. (2023). Context-specific regulation of extracellular vesicle biogenesis and cargo selection. Nat. Rev. Mol. Cell Biol. 24, 454–476. 10.1038/s41580-023-00576-0.

43. Lee, R.S., Monigatti, F., Briscoe, A.C., Waldon, Z., Freeman, M.R., and Steen, H. (2008). Optimizing sample handling for urinary proteomics. J. Proteome Res. 7, 4022–4030. 10.1021/pr800301h.

44. Thomas, C.E., Sexton, W., Benson, K., Sutphen, R., and Koomen, J. (2010). Urine collection and processing for protein biomarker discovery and quantification. Cancer Epidemiol. Biomark. Prev. Publ. Am. Assoc. Cancer Res. Cosponsored Am. Soc. Prev. Oncol. 19, 953–959. 10.1158/1055-9965.EPI-10-0069.

45. Erozenci, L.A., Pham, T.V., Piersma, S.R., Dits, N.F.J., Jenster, G.W., van Royen, M.E., Moorselaar, R.J.A., Jimenez, C.R., and Bijnsdorp, I.V. (2021). Simple urine storage protocol for extracellular vesicle proteomics compatible with at-home self-sampling. Sci. Rep. 11, 20760. 10.1038/s41598-021-00289-4.

46. Oeyen, E., Willems, H., ’t Kindt, R., Sandra, K., Boonen, K., Hoekx, L., De Wachter, S., Ameye, F., and Mertens, I. (2019). Determination of variability due to biological and technical variation in urinary extracellular vesicles as a crucial step in biomarker discovery studies. J. Extracell. Vesicles 8, 1676035. 10.1080/20013078.2019.1676035.

47. Rabas, N., Palmer, S., Mitchell, L., Ismail, S., Gohlke, A., Riley, J.S., Tait, S.W.G., Gammage, P., Soares, L.L., Macpherson, I.R., et al. (2021). PINK1 drives production of mtDNA-containing extracellular vesicles to promote invasiveness. J. Cell Biol. 220, e202006049. 10.1083/jcb.202006049.

48. McLelland, G.-L., Soubannier, V., Chen, C.X., McBride, H.M., and Fon, E.A. (2014). Parkin and PINK1 function in a vesicular trafficking pathway regulating mitochondrial quality control. EMBO J. 33, 282–295. 10.1002/embj.201385902.

49. Otto, J.J., Correll, V.L., Engstroem, H.A., Hitefield, N.L., Main, B.P., Albracht, B., Johnson-Pais, T., Yang, L.F., Liss, M., Boutros, P.C., et al. (2020). Targeted Mass Spectrometry of a Clinically Relevant PSA Variant from Post-DRE Urines for Quantitation and Genotype Determination. Proteomics Clin. Appl. 14, e2000012. 10.1002/prca.202000012.

50. Clayton, A., Court, J., Navabi, H., Adams, M., Mason, M.D., Hobot, J.A., Newman, G.R., and Jasani, B. (2001). Analysis of antigen presenting cell derived exosomes, based on immuno-magnetic isolation and flow cytometry. J. Immunol. Methods 247, 163–174. 10.1016/s0022-1759(00)00321-5.

51. Kim, Y., Jeon, J., Mejia, S., Yao, C.Q., Ignatchenko, V., Nyalwidhe, J.O., Gramolini, A.O., Lance, R.S., Troyer, D.A., Drake, R.R., et al. (2016). Targeted proteomics identifies liquid-biopsy signatures for extracapsular prostate cancer. Nat. Commun. 7, 11906. 10.1038/ncomms11906.

52. EV-TRACK Consortium, Van Deun, J., Mestdagh, P., Agostinis, P., Akay, Ö., Anand, S., Anckaert, J., Martinez, Z.A., Baetens, T., Beghein, E., et al. (2017). EV-TRACK: transparent reporting and centralizing knowledge in extracellular vesicle research. Nat. Methods 14, 228–232. 10.1038/nmeth.4185.

53. Human Protein Atlas proteinatlas.org.

54. Uhlén, M., Karlsson, M.J., Hober, A., Svensson, A.-S., Scheffel, J., Kotol, D., Zhong, W., Tebani, A., Strandberg, L., Edfors, F., et al. (2019). The human secretome. Sci. Signal. 12, eaaz0274. 10.1126/scisignal.aaz0274.

55. Thul, P.J., Åkesson, L., Wiking, M., Mahdessian, D., Geladaki, A., Ait Blal, H., Alm, T., Asplund, A., Björk, L., Breckels, L.M., et al. (2017). A subcellular map of the human proteome. Science 356, eaal3321. 10.1126/science.aal3321.

56. Berger, S.T., Ahmed, S., Muntel, J., Cuevas Polo, N., Bachur, R., Kentsis, A., Steen, J., and Steen, H. (2015). MStern Blotting-High Throughput Polyvinylidene Fluoride (PVDF) Membrane-Based Proteomic Sample Preparation for 96-Well Plates. Mol. Cell. Proteomics MCP 14, 2814–2823. 10.1074/mcp.O115.049650.

57. Kulak, N.A., Pichler, G., Paron, I., Nagaraj, N., and Mann, M. (2014). Minimal, encapsulated proteomic-sample processing applied to copy-number estimation in eukaryotic cells. Nat. Methods 11, 319–324. 10.1038/nmeth.2834.

58. Cox, J., Hein, M.Y., Luber, C.A., Paron, I., Nagaraj, N., and Mann, M. (2014). Accurate proteome-wide label-free quantification by delayed normalization and maximal peptide ratio extraction, termed MaxLFQ. Mol. Cell. Proteomics MCP 13, 2513–2526. 10.1074/mcp.M113.031591.

59. Fermin, D., Basrur, V., Yocum, A.K., and Nesvizhskii, A.I. (2011). Abacus: a computational tool for extracting and pre-processing spectral count data for label-free quantitative proteomic analysis. Proteomics 11, 1340–1345. 10.1002/pmic.201000650.

60. Schwanhäusser, B., Busse, D., Li, N., Dittmar, G., Schuchhardt, J., Wolf, J., Chen, W., and Selbach, M. (2011). Global quantification of mammalian gene expression control. Nature 473, 337–342. 10.1038/nature10098.

61. Tyanova, S., Temu, T., Sinitcyn, P., Carlson, A., Hein, M.Y., Geiger, T., Mann, M., and Cox, J. (2016). The Perseus computational platform for comprehensive analysis of (prote)omics data. Nat. Methods 13, 731–740. 10.1038/nmeth.3901.

62. Hänzelmann, S., Castelo, R., and Guinney, J. (2013). GSVA: gene set variation analysis for microarray and RNA-seq data. BMC Bioinformatics 14, 7. 10.1186/1471-2105-14-7.

63. Théry, C., Witwer, K.W., Aikawa, E., Alcaraz, M.J., Anderson, J.D., Andriantsitohaina, R., Antoniou, A., Arab, T., Archer, F., Atkin-Smith, G.K., et al. (2018). Minimal information for studies of extracellular vesicles 2018 (MISEV2018): a position statement of the International Society for Extracellular Vesicles and update of the MISEV2014 guidelines. J. Extracell. Vesicles 7, 1535750. 10.1080/20013078.2018.1535750.

64. Szklarczyk, D., Kirsch, R., Koutrouli, M., Nastou, K., Mehryary, F., Hachilif, R., Gable, A.L., Fang, T., Doncheva, N.T., Pyysalo, S., et al. (2023). The STRING database in 2023: protein-protein association networks and functional enrichment analyses for any sequenced genome of interest. Nucleic Acids Res. 51, D638–D646. 10.1093/nar/gkac1000.

